# Structure-activity relationship of pyrrolidine pentamine derivatives as inhibitors of the aminoglycoside 6′-*N*-acetyltransferase type Ib

**DOI:** 10.1101/2024.05.14.594018

**Authors:** Jan Sklenicka, Tung Tran, Maria S. Ramirez, Haley M. Donow, Angel J. Magaña, Travis LaVoi, Yasir Mamun, Prem Chapagain, Radleigh Santos, Clemencia Pinilla, Marc A. Giulianotti, Marcelo E. Tolmasky

## Abstract

Resistance to amikacin and other major aminoglycosides is commonly due to enzymatic acetylation by aminoglycoside 6′-*N* -acetyltransferase type I enzyme, of which type Ib [AAC(6′)-Ib] is the most widespread among Gram-negative pathogens. Finding enzymatic inhibitors could be an effective way to overcome resistance and extend the useful life of amikacin. Small molecules possess multiple properties that make them attractive compounds to be developed as drugs. Mixture-based combinatorial libraries and positional scanning strategy led to the identification of a chemical scaffold, pyrrolidine pentamine, that, when substituted with the appropriate functionalities at five locations (R1 - R5), inhibits AAC(6′)-Ib-mediated inactivation of amikacin. Structure-activity relationship (SAR) studies showed that while truncations to the molecule result in loss of inhibitory activity, modifications of functionalities and stereochemistry have different effects on the inhibitory properties. In this study, we show that alterations at position R1 of the two most active compounds, **2700.001** and **2700.003**, reduced inhibition levels, demonstrating the essential nature not only of the presence of an *S* -phenyl moiety at this location but also the distance to the scaffold. On the other hand, modifications on the R3, R4, and R5 positions have varied effects, demonstrating the potential for optimization. A correlation analysis between molecular docking values (ΔG) and the dose required for two-fold potentiation of compounds described in this and the previous studies showed a significant correlation between ΔG values and inhibitory activity.

**Highlights:**

- Amikacin resistance in Gram-negatives is mostly caused by the AAC(6′)-Ib enzyme
- AAC(6′)-Ib has been identified in most Gram-negative pathogens
- Inhibitors of AAC(6′)-Ib could be used to treat resistant infections
- Combinatorial libraries and positional scanning identified an inhibitor
- The lead compound can be optimized by structure activity relationship studies

## 1. Introduction

Multidrug resistance is one of the top concerns for human health. A growing number of people are dying due to acquiring resistant infections. At the same time, the multidrug resistance crisis is causing the cost of treatment to skyrocket [1–3], further compounding the issue. The number of new antimicrobials being developed falls short of what would be necessary for effectively managing the problem [4, 5]. Furthermore, only one of the recently approved antibiotics, cefiderocol, can be used against a bacterium included in the WHO list of critical pathogens [4]. Therefore, repurposing or extending the useful life of existing antimicrobials is essential to increase the armamentarium against the growing number of multidrug resistance bacterial pathogens [6]. Aminoglycoside antibiotics have been an instrumental component of the armamentarium in treating life-threatening infections [7, 8]. However, their spectrum of action is being diminished by the rise in resistance, mainly due to enzymatic modification catalyzed by aminoglycoside modifying enzymes (AMEs) [7, 9–11]. Significant efforts focused on designing new semisynthetic aminoglycosides by altering those found in nature to produce molecules refractory to the action AMEs [10, 12]. While these efforts resulted in the introduction of novel aminoglycosides like amikacin or plazomicin, attempts to identify or design inhibitors of the inactivating action of AMEs have been limited [6, 10, 11, 13, 14]. Consequently, no inhibitor has yet been introduced at the clinical level. Finding one suitable for human use will permit the design of combination therapies that are effective against resistant bacteria, thus extending the useful life and scope of existing aminoglycosides [6].

Amikacin is an aminoglycoside of high clinical relevance, but resistance—usually caused by the action of the aminoglycoside 6′-*N* -acetyltransferase type Ib [AAC(6′)-Ib]— abound in numerous geographical regions [10, 11, 15, 16]. Recent efforts to produce inhibitors of resistance mediated by this enzyme include exploring antisense strategies to turn off the expression of the *aac(6′)-Ib* gene and the identification of various chemicals that interfere with the acetylation reaction [13, 17–25]. In particular, small molecule inhibitors have the potential to serve as inhibitors of enzyme-mediated antibiotic resistance [26–28]. Developing an inhibitor that can be combined with amikacin could be an option to treat infections caused by multidrug-resistant strains that can no longer be controlled by carbapenems and other antimicrobials [29].

A recent study using mixture-based combinatorial libraries and the positional scanning strategy [30] identified a substituted pyrrolidine pentamine as a promising inhibitor of AAC(6′)-Ib-mediated acetylation of amikacin and other aminoglycosides [23] (Fig. 1A and Table 1). However, recent work revealed that some bacterial strains produce AAC(6′)-Ib in quantities far exceeding those needed for clinical resistance [16, 31]. This finding highlights the necessity for an inhibitor with enough potency to effectively combat a range of pathogenic bacteria, even in environments with high enzyme concentrations. In addressing this challenge, we conducted further structure-activity relationship (SAR) studies to investigate how changes in the compounds’ stereochemistry and substitutions of functionalities impact their inhibitory effectiveness [24]. Expanding upon our previous SAR analyses, this work aims to continue exploring the connection between molecular modifications and their inhibitory effects. Such insights are instrumental for designing a potent inhibitor that, in combination with amikacin, could overcome the resistance conferred by AAC(6′)-Ib.

**Figure 1.**
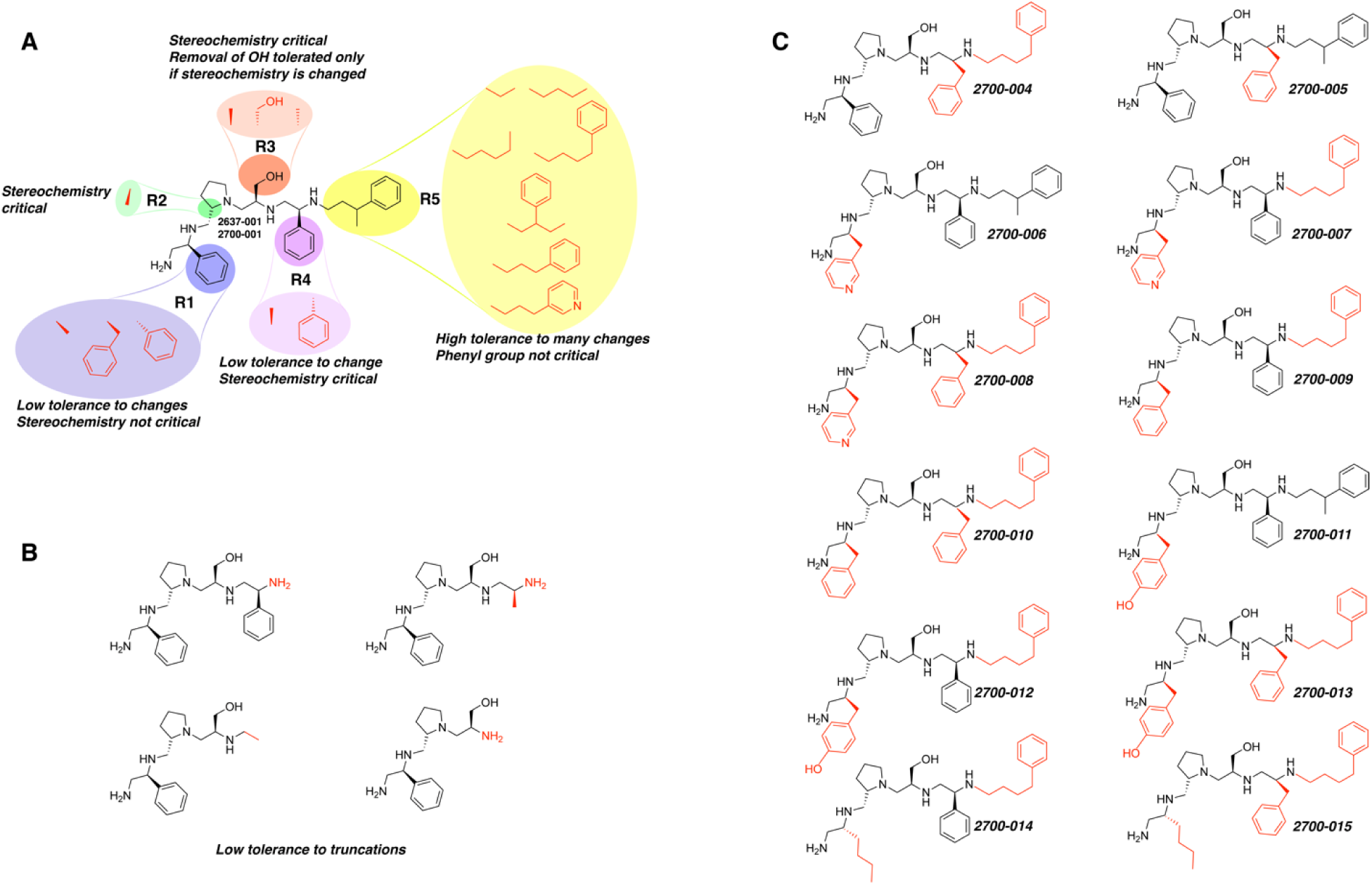
Compounds structures. A. Chemical structure of compound **2700.001** (formerly **2637-001**) showing the pyrrolidine pentamine scaffold with the two S-phenyl groups (blue and purple background), the S-hydroxymethyl group (orange background), and the 3-phenylbutyl group (yellow background) at the positions R1, R3, R4, and R5, respectively. Substitutions previously evaluated are shown in red over the corresponding color background. B. Compounds resulting from truncations of various sizes to **2700.001**. C. Chemical structures of compounds assessed in this work. Substitutions with respect to **2700.001** are shown in red.

**Table 1.**
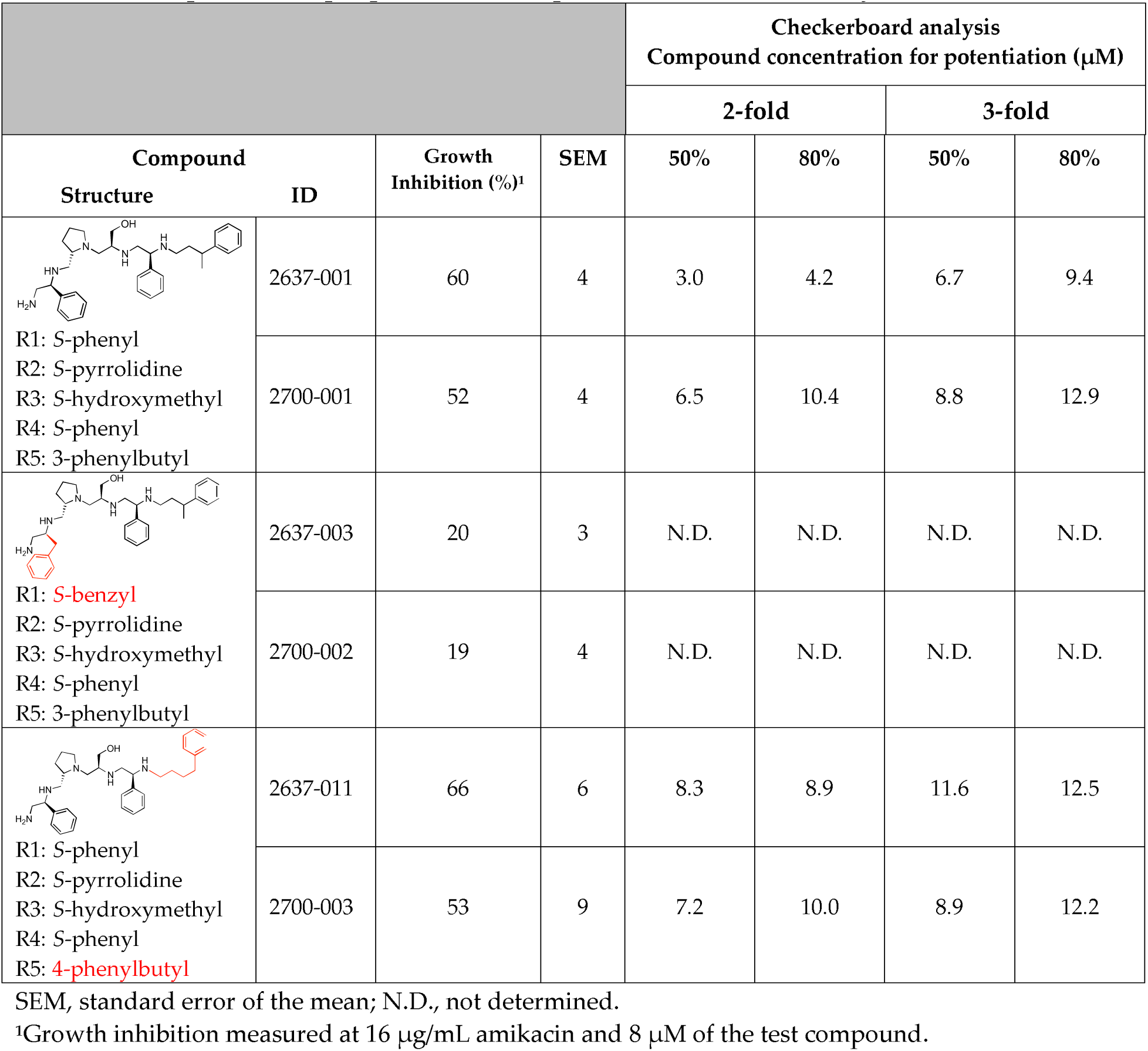
Comparison of properties of compounds from different synthesis batches.

|  |  |  |  | Checkerboard analysis |  |  |  |
| --- | --- | --- | --- | --- | --- | --- | --- |
|  |  |  |  | Compound concentration for potentiation (μM) |  |  |  |
| Compound |  | Growth Inhibition (%)<sup>1</sup> | SEM | 2-fold |  | 3-fold |  |
| Structure | ID |  |  | 50% | 80% | 50% | 80% |
| R1: S-phenyl<br>R2: S-pyrrolidine<br>R3: S-hydroxymethyl<br>R4: S-phenyl<br>R5: 3-phenylbutyl | 2637-001 | 60 | 4 | 3.0 | 4.2 | 6.7 | 9.4 |
|  | 2700-001 | 52 | 4 | 6.5 | 10.4 | 8.8 | 12.9 |
| R1: S-benzyl<br>R2: S-pyrrolidine<br>R3: S-hydroxymethyl<br>R4: S-phenyl<br>R5: 3-phenylbutyl | 2637-003 | 20 | 3 | N.D. | N.D. | N.D. | N.D. |
|  | 2700-002 | 19 | 4 | N.D. | N.D. | N.D. | N.D. |
| R1: S-phenyl<br>R2: S-pyrrolidine<br>R3: S-hydroxymethyl<br>R4: S-phenyl<br>R5: 4-phenylbutyl | 2637-011 | 66 | 6 | 8.3 | 8.9 | 11.6 | 12.5 |
|  | 2700-003 | 53 | 9 | 7.2 | 10.0 | 8.9 | 12.2 |

SEM, standard error of the mean; N.D., not determined.
<sup>1</sup>Growth inhibition measured at 16 μg/mL amikacin and 8 μM of the test compound.

## 2. Materials and Methods

### 2.1. Bacterial Strains and Small Molecule Compounds

*A. baumannii* A155, a clonal complex 109 multidrug resistant strain that harbors *aac(6’)*, was isolated from a urinary sample at a hospital in the Autonomous City of Buenos Aires, Argentina [32–34]. Solid and liquid routine cultures were carried out in Lennox L broth (1% tryptone, 0.5% yeast extract, 0.5% NaCl) with the addition or not of 2% agar. Cultures to determine levels of amikacin resistance to amikacin were performed in Mueller-Hinton broth. The compounds were synthesized at the Center for Translational Science at Florida International University, as described previously [23]. Briefly, a polyamide scaffold was synthesized on a solid support using standard Boc-protected chemistry. Then, the amide residues were reduced with borane, and the compounds were removed from the solid support using hydrofluoric acid (Fig. S1, Supplementary material). The purity and identity of compounds were verified as before [24] using a Shimadzu 2020 Liquid chromatography-mass spectrometry (LCMS) system. Chromatographic separations were carried out on a Phenomenex Luna C18 analytical column (5 μm, 150 mm × 4.6 mm i.d.) with a Phenomenex C18 column guard (5 μm, 4 × 3.0 mm i.d.). The equipment was controlled and integrated with the Shimadzu LCMS solutions software version 5. The mobile phases A and B for LCMS analysis were LCMS-grade water and LCMS-grade acetonitrile, respectively (Sigma-Aldrich and Fisher Scientific, both with 0.1% formic acid for a pH of 2.7). The procedure for analyzing 5 μL aliquots was identical to that described in our previous study [24]. Figures S2 and S3 (Supplementary material) show the relevant information on the compounds’ characterization and degree of purification.

### 2.2. Initial Growth Inhibition Assays

An initial test to determine the levels of inhibition produced by the compounds was done by measuring OD_600_ after 20 h of growth in Mueller-Hinton broth supplemented with 16 μg/ml amikacin and 8 μM of the potential inhibitor. These concentrations were selected based on previous studies using compounds with the same scaffold [23, 24]. Each compound was tested in four separate experiments by duplicate. The data values, expressed in percent inhibition based on the OD_600_ measurements, were adjusted using the mixture-modeling described below to account for compound inhibition and averaged. The standard error of the mean for n = 8 percentage growth inhibition values of each compound was calculated. Each *p*-value for testing the difference in inhibitory activity between a compound and **2700.001** was calculated using a two-sample t-test with Bonferroni– Holm correction. A *p*-value of less than 0.05 was considered significant. All compounds that did not show a substantial reduction in inhibitory activity with respect to compound **2700.001** were used in checkerboard assays.

### 2.3. Checkerboard Assays

Checkerboard assays were performed in Mueller-Hinton broth as described previously [23, 24]. Each compound was tested on at least three independent checkerboard experiments, in which each dose combination was tested by duplicate. The variables were the potential inhibitor (tested at 0, 4, 8, 16, and 24 μM) and amikacin (tested at 0, 8, 16, 32, and 64 μg/mL). Assays were carried out in microtiter plates using the BioTek Synergy 5 microplate reader (BioTek Synergy 5). The data were analyzed applying an approach that quantifies exact levels of synergy, i.e., eliminating any antibacterial effect exerted by the inhibitor [23, 24, 35]. The model considers that amikacin and the potential inhibitors have independent antibacterial mechanisms of action. The percent activity of the mixture of the two chemicals was modeled as follows:

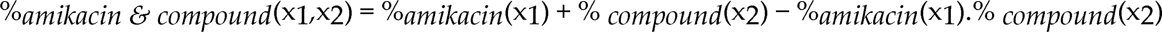

Where x_1_ and x_2_ are the amikacin and tested compound concentrations, respectively. The effective percent activity of the antibiotic alone at a given concentration, after accounting for compound activity, can be calculated using a rearrangement of the previous equation as follows:

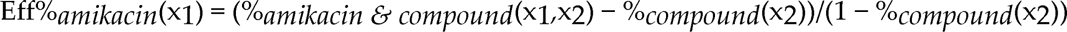

These calculations provide the actual change in amikacin resistance levels. The above methodology was applied to the median of the values at each dose combination. Once applied to the checkerboard data, the mean effective concentration of amikacin to achieve 50% and 80% inhibition (IC_50_/IC_80_) at each dose of the potentiating compound was determined using pairwise interpolation. The fold potentiation of amikacin for each dose of the compounds and each dose point was then calculated. Finally, the compound concentration needed to achieve 2- and 3-fold potentiation for each of the two dose points was determined using pairwise interpolation.

### 2.4. Molecular Docking

To prepare the receptor for docking experiments, the x-ray crystal structure of AAC(6′)-Ib complexed with kanamycin C and AcetylCoA [36] was obtained from the protein data bank (PDB 1V0C). Kanamycin C was removed from the AAC(6′)-Ib protein structure, and the final structure was converted to pdbqt format using AutoDockTools 4.2 [37]. The cavity where kanamycin C was bound to the protein was selected as the target site for virtual screening. Again, using AutoDockTools 4.2, residues W49, Y65, E73, V75, Q91, Y93, S98, D100, W103, D115, D152, and D179 were converted to flexible residues to allow for flexible binding. Next, the ligands were prepared by converting the structures of the compounds to 3D with polar hydrogen bonds and finally in pdbqt format using Open Babel [38]. Molecular docking and screening were performed using AutoDock Vina 1.2 [39]. The docking scores were sorted and ranked based on their predicted binding energies (delta G, Kcal/mol), with the lowest score representing the best binding. LigPlot+ [40] was used to generate a 2D ligand-protein interaction map. PyMol 2.3 [41] was used for visualization and rendering.

## 3. Results

The high relevance of AAC(6′)-Ib as the cause of resistance to amikacin in pathogenic Gram-negatives motivated the search for inhibitors of the enzymatic acetylation that inactivates the aminoglycoside molecule. Utilizing mixture-based combinatorial libraries and the positional scanning strategy, we identified compound **2637.001,** which consists of a pyrrolidine pentamine scaffold with two *S* -phenyl groups, an *S* -hydroxymethyl group, and a 3-phenylbutyl group at positions R1, R3, R4, and R5, respectively, as shown in Figure 1A. To study the potential interactions of compound **2637.001** with the AAC(6′)-Ib molecule and its inhibitory activity, a series of compound **2637.001** analogs were analyzed. Figures 1A and 1B graphically show the tolerance and effects of substituting the chemical groups at each location, modifying the stereochemical conformation at R2, or reducing the size of the molecule. Position R1 showed a low tolerance to modifications, including changing the stereochemistry, replacing the phenyl with a methyl group, and increasing the distance between the phenyl group and the scaffold.

Position R4 exhibited low tolerance to modifying the stereochemistry or replacing the phenyl with a methyl group. To gain further insights into the contribution or effect of substitutions at these positions in combination with substitutions at the R5 position, which has high tolerance to modifications, we generated a collection of twelve analogs with modifications at positions R1, R4, and R5 (Fig. 1C, identified as **2700** series). The R1 location was unmodified or modified to carry a heteroatom in the aromatic moiety, to increase the distance between the phenyl or pyridine groups and the scaffold, or to replace the aromatic moiety with an aliphatic one that occupies approximately the same space. The R4 position was unmodified or modified to move the phenyl group away from the scaffold by inserting a methylene moiety. The R5 position was unmodified or modified by replacing the 3-phenylbutyl with a 4-phenylbutyl group.

Each compound’s efficacy as an amikacin resistance inhibitor was evaluated on *A. baumannii* A155, a strain harboring the *aac(6*′*)-Ib* gene, at concentrations of 16 μg/mL amikacin and 8 μM of the test compound. Previous results showed that *A. baumannii* A155 can grow in 16 μg/mL amikacin containing Mueller-Hinton broth [24]. The growth curves of the cultures were determined by measuring OD_600_ every 20 m, and the values after 20 h incubation, when the cultures were already in stationary phase, were used to calculate the percentage of growth inhibition with respect to bacteria growing in media with the sole addition of amikacin.

To ensure that the results obtained with compounds synthesized at different times could be compared, an experiment was carried out to determine the inhibition levels of three compounds synthesized first for the previous study [24] and then for the present study (**2637.001,** now **2700.001**; **2637.003,** now **2700.002**; **2637.011,** now called **2700.003)**. Comparisons were carried out at a single dose and using checkerboard assays for higher accuracy in the case of the two structures that showed more potent inhibition of amikacin resistance. The results of these tests are shown in Table 1. The inhibition levels observed when using compounds from both series were sufficiently close to confirm that the results obtained with the most recently designed compounds can be compared to the previous analyses.

Considering the features of the two compounds with higher inhibiting activity (**2700.001** and **2700.003**), a collection of analogs was generated to gain deeper insights into the effect of modifications at different positions. Inspection of the inhibition of amikacin resistance exerted by these compounds showed that one out of the twelve newly designed analogs, **2700.004,** appeared to restore resistance to amikacin at levels comparable to compounds **2700.001** and **2700.003** (Table 2).

**Table 2.**
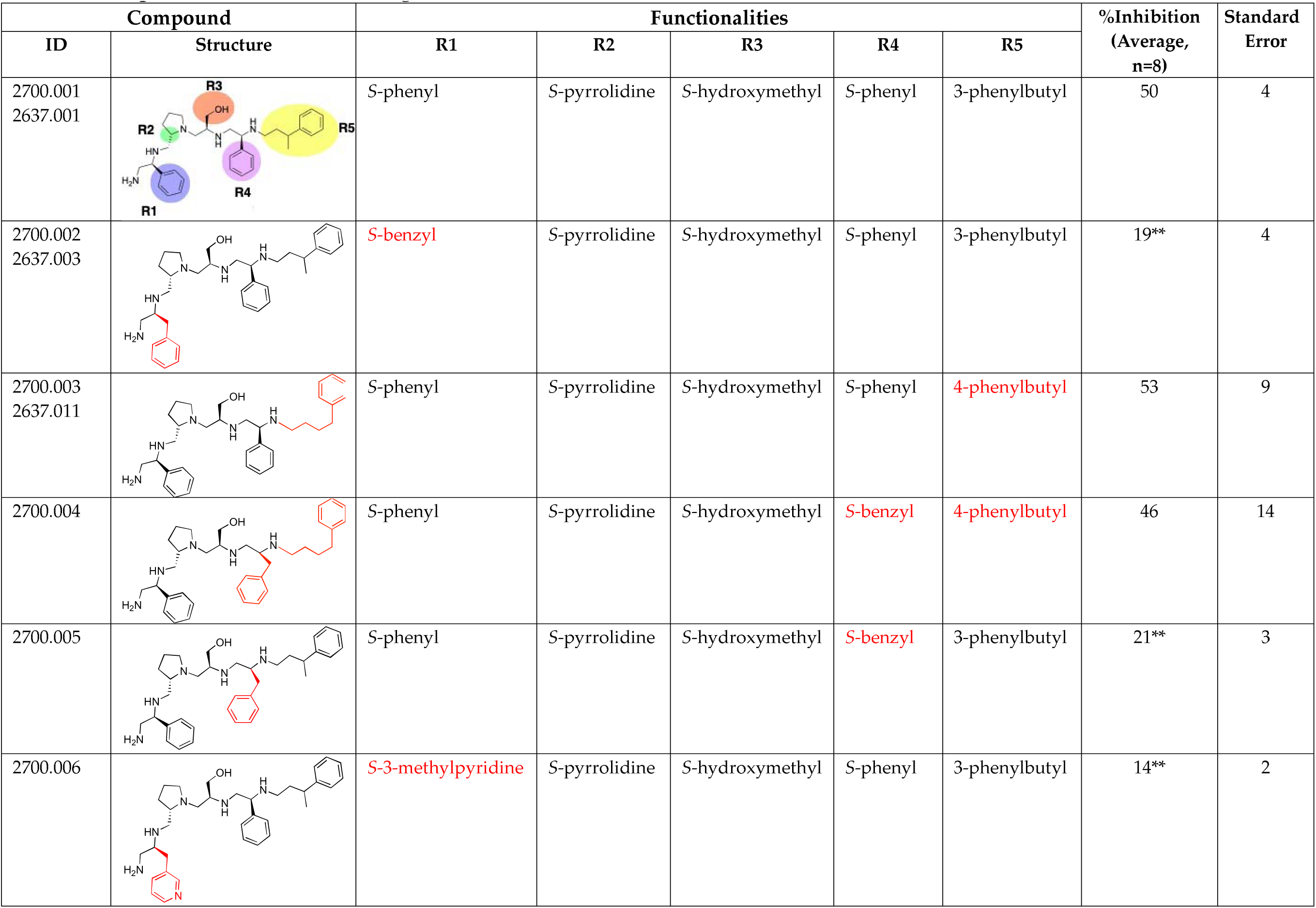
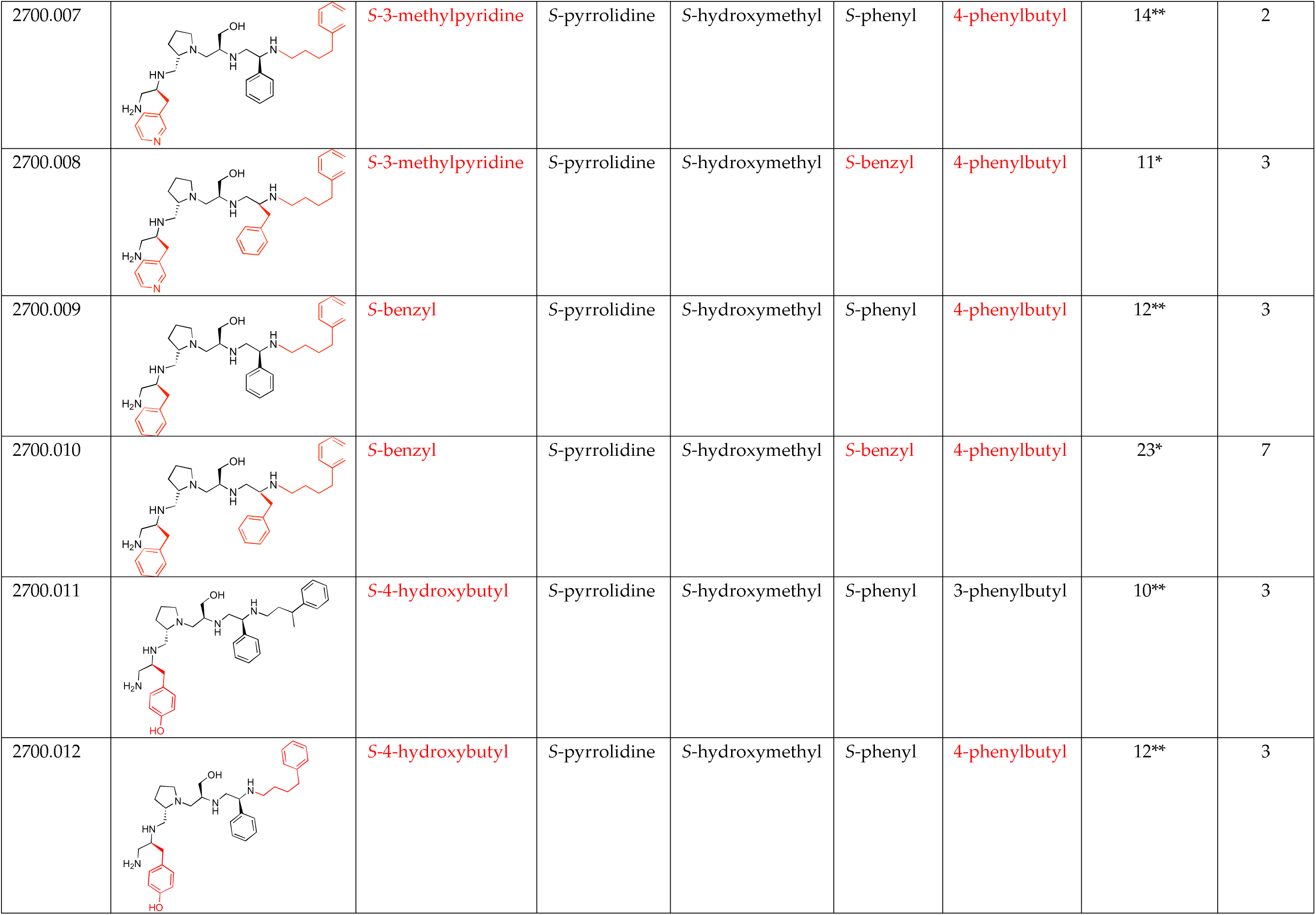
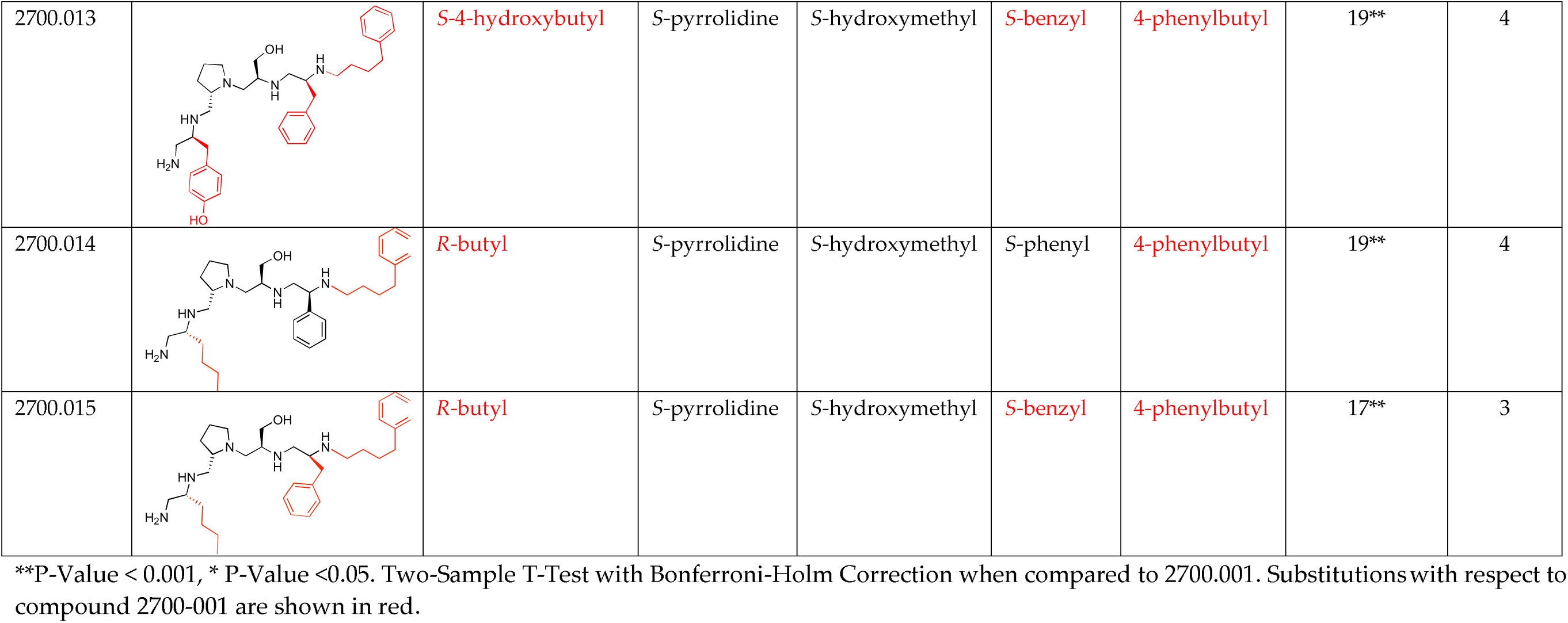
Properties of 2700.001 analogs.

| Compound |  | Functionalities |  |  |  |  | %Inhibition<br>(Average,<br>n=8) | Standard<br>Error |
| --- | --- | --- | --- | --- | --- | --- | --- | --- |
| ID | Structure | R1 | R2 | R3 | R4 | R5 |  |  |
| 2700.001<br>2637.001 |  | S-phenyl | S-pyrrolidine | S-hydroxymethyl | S-phenyl | 3-phenylbutyl | 50 | 4 |
| 2700.002<br>2637.003 |  | S-benzyl | S-pyrrolidine | S-hydroxymethyl | S-phenyl | 3-phenylbutyl | 19** | 4 |
| 2700.003<br>2637.011 |  | S-phenyl | S-pyrrolidine | S-hydroxymethyl | S-phenyl | 4-phenylbutyl | 53 | 9 |
| 2700.004 |  | S-phenyl | S-pyrrolidine | S-hydroxymethyl | S-benzyl | 4-phenylbutyl | 46 | 14 |
| 2700.005 |  | S-phenyl | S-pyrrolidine | S-hydroxymethyl | S-benzyl | 3-phenylbutyl | 21** | 3 |
| 2700.006 |  | S-3-methylpyridine | S-pyrrolidine | S-hydroxymethyl | S-phenyl | 3-phenylbutyl | 14** | 2 |
| 2700.007 |  | S-3-methylpyridine | S-pyrrolidine | S-hydroxymethyl | S-phenyl | 4-phenylbutyl | 14** | 2 |
| 2700.008 |  | S-3-methylpyridine | S-pyrrolidine | S-hydroxymethyl | S-benzyl | 4-phenylbutyl | 11* | 3 |
| 2700.009 |  | S-benzyl | S-pyrrolidine | S-hydroxymethyl | S-phenyl | 4-phenylbutyl | 12** | 3 |
| 2700.010 |  | S-benzyl | S-pyrrolidine | S-hydroxymethyl | S-benzyl | 4-phenylbutyl | 23* | 7 |
| 2700.011 |  | S-4-hydroxybutyl | S-pyrrolidine | S-hydroxymethyl | S-phenyl | 3-phenylbutyl | 10** | 3 |
| 2700.012 |  | S-4-hydroxybutyl | S-pyrrolidine | S-hydroxymethyl | S-phenyl | 4-phenylbutyl | 12** | 3 |
| 2700.013 |  | *S*-4-hydroxybutyl | *S*-pyrrolidine | *S*-hydroxymethyl | *S*-benzyl | 4-phenylbutyl | 19** | 4 |
| 2700.014 |  | *R*-butyl | *S*-pyrrolidine | *S*-hydroxymethyl | *S*-phenyl | 4-phenylbutyl | 19** | 4 |
| 2700.015 |  | *R*-butyl | *S*-pyrrolidine | *S*-hydroxymethyl | *S*-benzyl | 4-phenylbutyl | 17** | 3 |

\*\*P-Value < 0.001, \* P-Value <0.05. Two-Sample T-Test with Bonferroni-Holm Correction when compared to 2700.001. Substitutions with respect to compound 2700-001 are shown in red.

The effect of altering compound **2700.001** by making a single substitution at the R1 position was assessed. Subsequently, three analogs consisting of moving the aromatic ring further from the scaffold (**2700.002**), replacing the aromatic ring with one containing a heteroatom (**2700.006**), or by a hydroxy-substituted one (**2700.011**) were evaluated. All three compounds with a single substitution at the R1 position showed reduced inhibition levels, demonstrating the importance of the *S* -phenyl moiety in the context of compound **2700.001**. Single substitutions at the R1 position were also assessed utilizing **2700.003** as a starting point. All these R1 substitutions, **2700.007** (the aromatic ring replaced with one containing a heteroatom), **2700.009 (**the aromatic ring moved further from the scaffold by inserting a methylene group), **2700.012 (**the aromatic ring replaced by a hydroxy-substituted one), and **2700.014** (the aromatic ring replaced by a linear aliphatic group) (Table 2) resulted in a significant reduction in inhibition levels, further confirming the importance of the S-phenyl moiety in the R1 position for our lead compounds.

The effect of making a single substitution at the R4 position of **2700.001** was assessed moving the phenyl moiety one carbon away from the scaffold. This modification significantly reduced inhibition levels (**2700.005**) (Table 2). Of note, this modification was well tolerated if made on the **2700.003** compound, when position R5 was occupied by a 4-phenylbutyl instead of the 3-phenylbutyl group present in compound **2700.004** (Table 2). Thus, the inhibitory effect of R1 single substitution of **2700.004** was also assessed. These four analog compounds, **2700.008**, **2700.010**, **2700.013**, and **2700.015,** showed significantly reduced inhibition compared to **2700.004** (Table 2). The results obtained using this set of analogs, taken together with those generated using the previous set [24], provide us insights into the sensitivity of R group substitutions and indicate the potential for further optimization with the R4 substitution of the **2700.001** and **2700.003** in future studies.

To assess the potentiating effect of amikacin by the compounds under investigation with higher accuracy, we selected those that produced inhibition levels higher than 20% to carry out checkerboard experiments. Additionally, two compounds that inhibited at less than 20% were selected as controls (**2700.007** and **2700.013**). Then, compounds **2700.001**, **2700.003**, **2700.004**, **2700.005**, **2700.007**, **2700.010**, and **2700.013** were tested in checkerboard assays carried out at 0, 4, 8, 16, and 32 μg/ml amikacin and 0, 2, 4, 8, 16, and 24 μM compound. The raw experimental values obtained were adjusted using mixture modeling (described in the Materials and Methods section) to account for any compound’s antimicrobial contribution to growth inhibition. These values were used to calculate the concentration of potentiated amikacin to achieve 50% and 80% bacterial inhibition of growth, the fold potentiation (over amikacin alone) of each of these dose points associated with each compound at each concentration, and the compound concentration needed to achieve 2- or 3-fold potentiation of both the 50% and 80% inhibitory dose points (Table 3 and Fig. S4, Supplementary material). The checkerboard results indicate that the most potent inhibitor among the compounds with newly synthesized structures within the 2700 series is inferior to those already identified in the previous studies [23, 24]. This includes **2700.004**, for which single-point potentiation assay results did not adequately demonstrate a difference from **2700.001**.

**Table 3.** Summary of checkerboard assays.

| Compound ID | Inhibition (μM) |  |  |  |
| --- | --- | --- | --- | --- |
|  | 2-Fold |  | 3-Fold |  |
|  | 50% | 80% | 50% | 80% |
| 2700.001 | 6.5 | 10.4 | 8.8 | 12.9 |
| 2700.003 | 7.2 | 10.0 | 8.9 | 12.2 |
| 2700.004 | 19.3 | 23.4 | > 24 | > 24 |
| 2700.005 | 14.2 | 15.0 | 19.9 | 19.5 |
| 2700.007 | > 24 | > 24 | > 24 | > 24 |
| 2700.010 | 16.2 | 15.5 | 19.9 | 19.6 |
| 2700.013 | > 24 | > 24 | > 24 | > 24 |

Compound concentration needed to achieve 2- or 3-fold potentiation of the 50% and 80% inhibitory dose points.

A correlation study between molecular docking values and checkerboard potentiation was performed integrating the results of the previous structure-activity relationship study [24] and the compounds presented herein. The ΔG (Kcal/mol) values were determined for the compounds across both sets for which checkerboard experiments were carried out (Table 4). Figure 2 shows a regression analysis considering the ΔG values and the compound dose required for a two-fold potentiation as determined by the checkerboard assays. The results demonstrate a significant correlation between ΔG values and inhibitory activity (r = 0.76, p = 0.0004). The complex of compounds (**2700.001**, **2700.004**, **2700.007**, and **27700.013**) with AAC(6′)-Ib obtained from molecular docking are shown in Figure S5 (Supplementary material).

**Figure 2.**
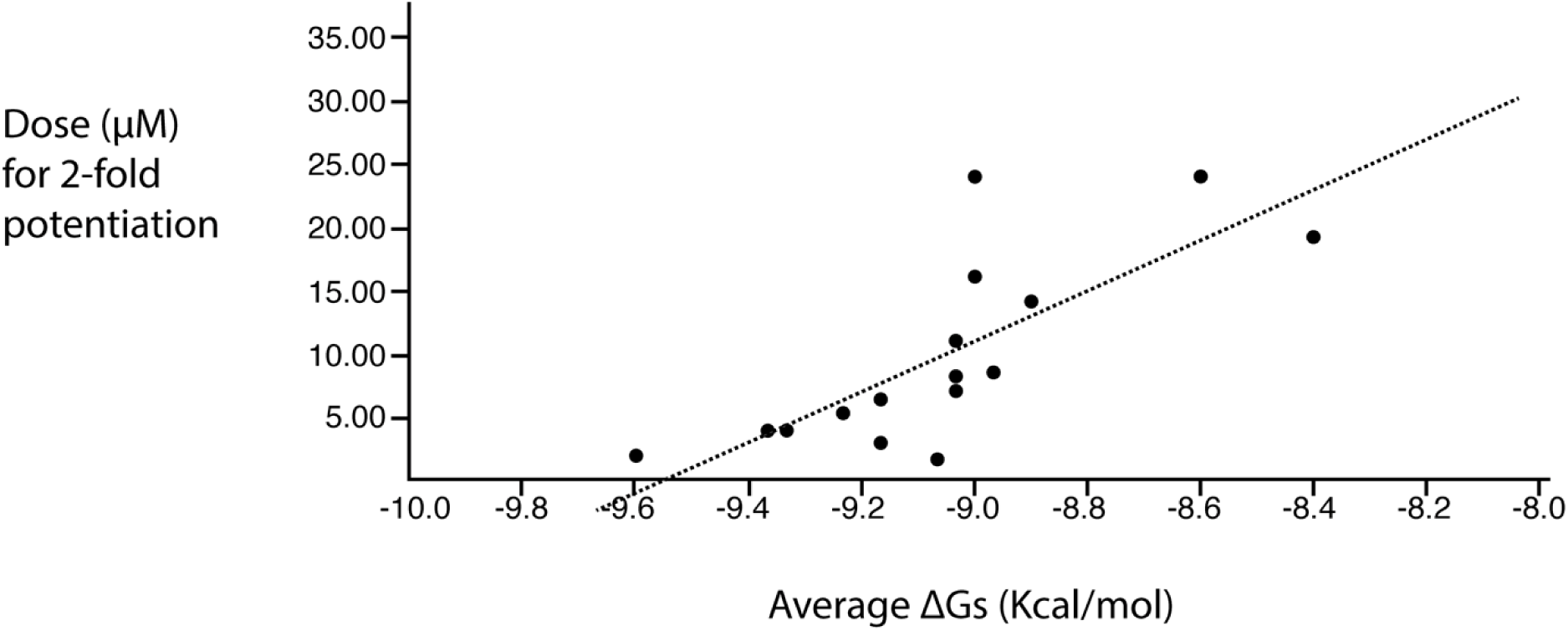
Predicted binding affinity vs experimentally measured potentiation of the different compounds. The x-axis shows the calculated ΔG average when the compound interacts with the enzyme molecule. The y-axis shows the calculated efficacy for potentiation of the compounds based on the experimental data. The regression line is included in the figure. The correlation is significant (r = 0.76; p = 0.0004). The values utilized are those shown in the two rightmost columns of Table 4.

**Table 4.**
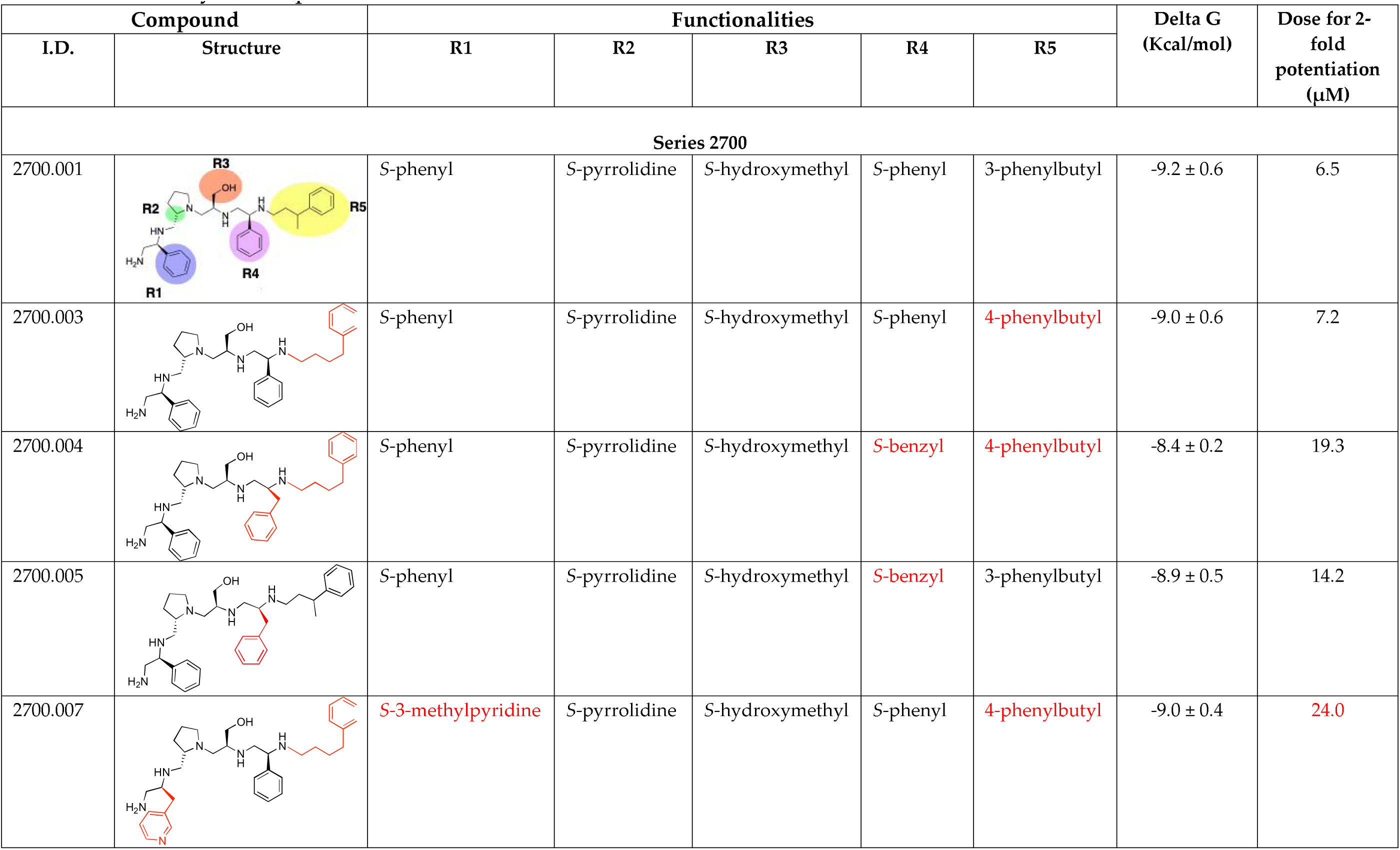
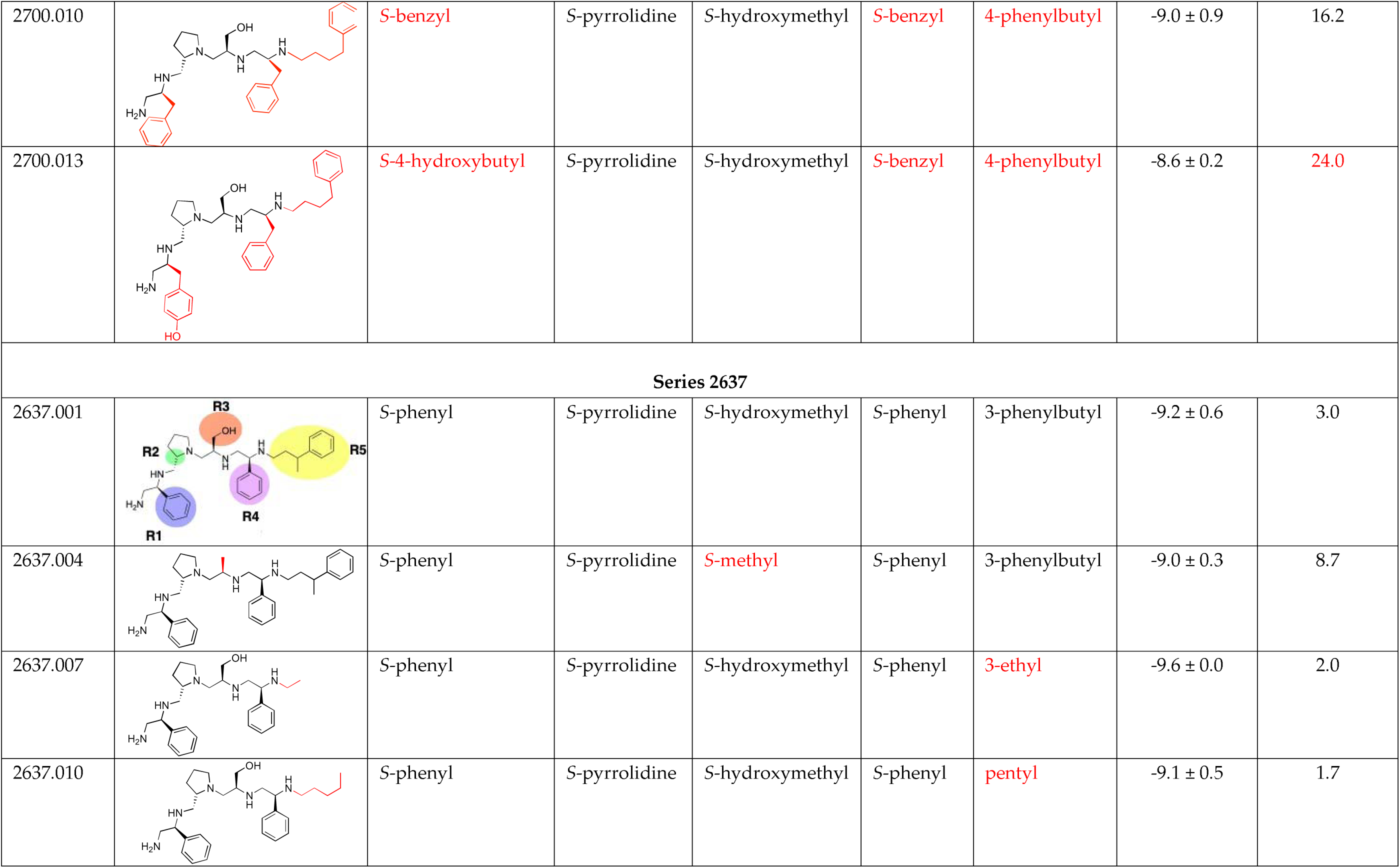
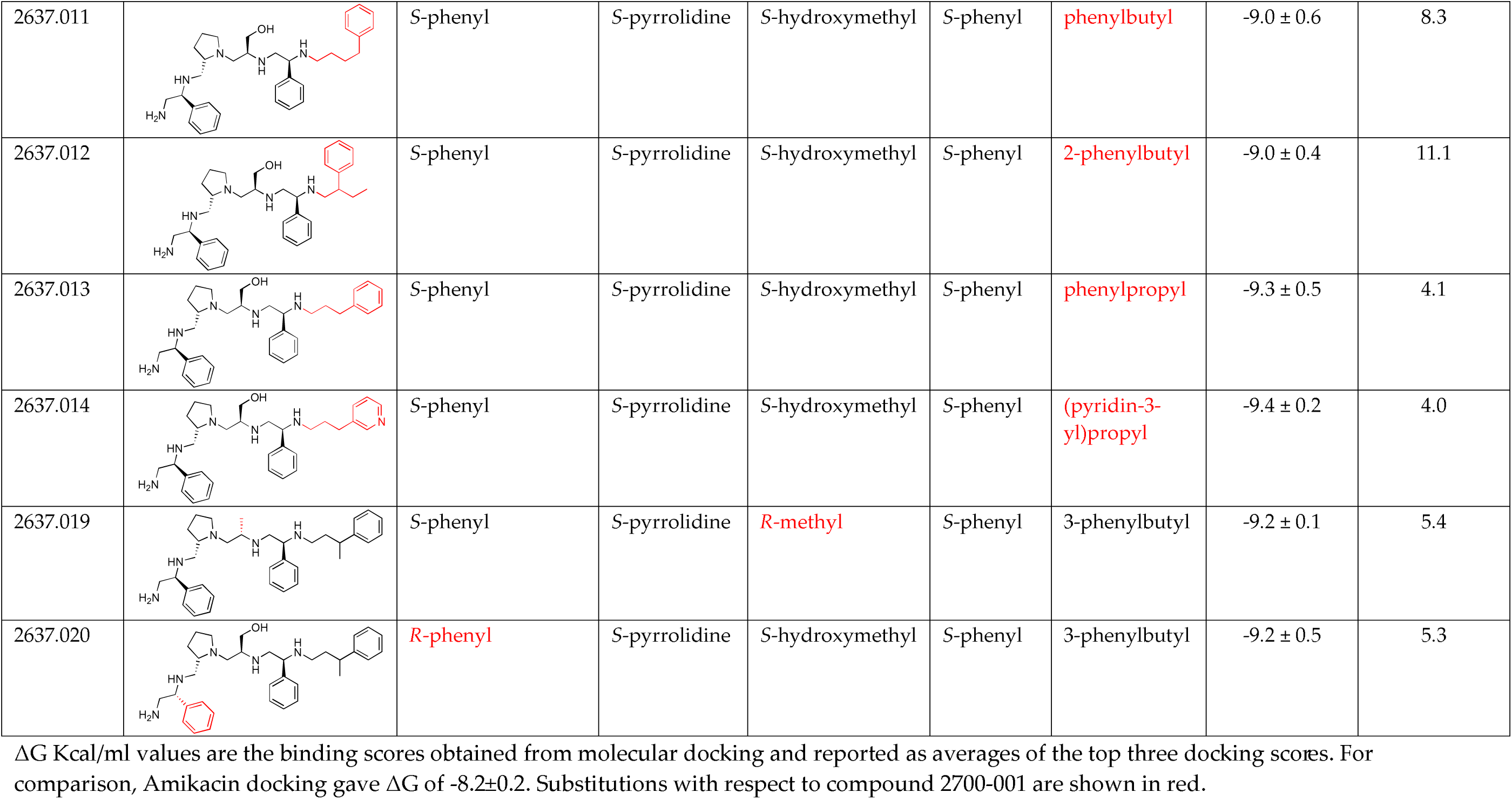
Summary of compounds tested on checkerboards.

| Compound |  | Functionalities |  |  |  |  | Delta G<br>(Kcal/mol) | Dose for 2-fold<br>potentiation<br>(μM) |
| --- | --- | --- | --- | --- | --- | --- | --- | --- |
| I.D. | Structure | R1 | R2 | R3 | R4 | R5 |  |  |
| Series 2700 |  |  |  |  |  |  |  |  |
| 2700.001 |  | S-phenyl | S-pyrrolidine | S-hydroxymethyl | S-phenyl | 3-phenylbutyl | -9.2 ± 0.6 | 6.5 |
| 2700.003 |  | S-phenyl | S-pyrrolidine | S-hydroxymethyl | S-phenyl | 4-phenylbutyl | -9.0 ± 0.6 | 7.2 |
| 2700.004 |  | S-phenyl | S-pyrrolidine | S-hydroxymethyl | S-benzyl | 4-phenylbutyl | -8.4 ± 0.2 | 19.3 |
| 2700.005 |  | S-phenyl | S-pyrrolidine | S-hydroxymethyl | S-benzyl | 3-phenylbutyl | -8.9 ± 0.5 | 14.2 |
| 2700.007 |  | S-3-methylpyridine | S-pyrrolidine | S-hydroxymethyl | S-phenyl | 4-phenylbutyl | -9.0 ± 0.4 | 24.0 |
| 2700.010 |  | S-benzyl | S-pyrrolidine | S-hydroxymethyl | S-benzyl | 4-phenylbutyl | -9.0 ± 0.9 | 16.2 |
| 2700.013 |  | S-4-hydroxybutyl | S-pyrrolidine | S-hydroxymethyl | S-benzyl | 4-phenylbutyl | -8.6 ± 0.2 | 24.0 |
| Series 2637 |  |  |  |  |  |  |  |  |
| 2637.001 |  | S-phenyl | S-pyrrolidine | S-hydroxymethyl | S-phenyl | 3-phenylbutyl | -9.2 ± 0.6 | 3.0 |
| 2637.004 |  | S-phenyl | S-pyrrolidine | S-methyl | S-phenyl | 3-phenylbutyl | -9.0 ± 0.3 | 8.7 |
| 2637.007 |  | S-phenyl | S-pyrrolidine | S-hydroxymethyl | S-phenyl | 3-ethyl | -9.6 ± 0.0 | 2.0 |
| 2637.010 |  | S-phenyl | S-pyrrolidine | S-hydroxymethyl | S-phenyl | pentyl | -9.1 ± 0.5 | 1.7 |
| 2637.011 |  | S-phenyl | S-pyrrolidine | S-hydroxymethyl | S-phenyl | phenylbutyl | -9.0 ± 0.6 | 8.3 |
| 2637.012 |  | S-phenyl | S-pyrrolidine | S-hydroxymethyl | S-phenyl | 2-phenylbutyl | -9.0 ± 0.4 | 11.1 |
| 2637.013 |  | S-phenyl | S-pyrrolidine | S-hydroxymethyl | S-phenyl | phenylpropyl | -9.3 ± 0.5 | 4.1 |
| 2637.014 |  | S-phenyl | S-pyrrolidine | S-hydroxymethyl | S-phenyl | (pyridin-3-yl)propyl | -9.4 ± 0.2 | 4.0 |
| 2637.019 |  | S-phenyl | S-pyrrolidine | R-methyl | S-phenyl | 3-phenylbutyl | -9.2 ± 0.1 | 5.4 |
| 2637.020 |  | R-phenyl | S-pyrrolidine | S-hydroxymethyl | S-phenyl | 3-phenylbutyl | -9.2 ± 0.5 | 5.3 |

$\Delta G$ Kcal/ml values are the binding scores obtained from molecular docking and reported as averages of the top three docking scores. For comparison, Amikacin docking gave $\Delta G$ of -8.2±0.2. Substitutions with respect to compound 2700-001 are shown in red.

The set of analogs analyzed in this study, in combination with the results obtained using the previous set [24], provides us insights into the sensitivity of R group substitutions. The data from this set of compounds highlights the importance of the R1 substitution of the original compound, confirms potential alterations for the R5 position, and indicates that there may still be opportunities to optimize the R3 and R4 substitutions of the **2700.001** and **2700.003** in future studies.

## 4. Discussion

Prolonging the life of available antibiotics is crucial. Designing or discovering adjuvants that inhibit the expression or activity of biomolecules responsible for resistance can help prevent or slow the emergence of untreatable infections [1, 6, 42]. Such infections could increase mortality rates not only from primary diseases but also by complicating medical and dental procedures [26]. Although this strategy has been extraordinarily successful in expanding the usefulness of β-lactams through the combination with β-lactamase inhibitors [43], it has not yet progressed beyond research settings for aminoglycosides [26]. Since the most widespread mechanism of resistance to aminoglycosides, including amikacin, involves acetylation catalyzed by the AAC(6′)- Ib enzyme [11, 16], finding inhibitors of this enzyme could allow for the treatment of numerous life-threatening infections, including those caused by carbapenem-resistant strains [21]. A proven methodology to identify bioactive compounds uses mixture-based combinatorial libraries and the positional scanning approach, which allows for identifying scaffold structures and testing large numbers of compounds simultaneously [23, 44, 45]. We recently identified a substituted pyrrolidine pentamine scaffold, compound **2700-001**, an inhibitor of the AAC(6′)-Ib enzymatic activity [23]. This compound was used as a starting point to introduce modifications at different locations along the molecule, as well as to remove portions of the molecule, as part of SAR studies. These analyses aimed to understand the interaction between the potential inhibitors and the enzyme, as well as to identify more potent inhibitors.

In a previous study testing a series of analogs, it was concluded that the integrity of the pyrrolidine pentamine scaffold and the stereochemistry at positions R2, R3, and R4 were necessary for the compounds’ ability to act as inhibitors of resistance to amikacin (Fig. 1, A and B)[24]. This analysis was expanded to explore the effects of additional modifications to the most active compounds identified to date (**2700.001** and **2700.003**). The new compounds were designed introducing one, two, or three substitutions at the R1, R4, or R5 positions (Fig. 1C and Table 1). A preliminary analysis comparing the same compounds from two independent synthesis events showed comparable results, thereby validating the findings and ensuring consistency across different synthesis batches and different biological activity determinations. Consistent with earlier results, a single substitution at the R1 position of compounds **2700.001** and **2700.003** led to a loss of inhibitory activity. Thus, the *S* -phenyl moiety remains the most effective substituent known to date for a pyrrolidine pentamine derivative to inhibit resistance. Substitutions at the R4 position of **2700.001** and **2700.003** resulted in loss of inhibitory activity with the only exception that replacing the *S*-phenyl group by *S* -benzyl in the compound **2700.003** produced a compound with activity comparable to that of compounds **2700.001** and **2700.003**. The results obtained with the latest group of analogs suggest that it may be possible to increase the inhibitory activity optimizing the R4 and R5 substitutions of the **2700.001** and **2700.003** compounds. Since the purpose of the single dose analysis was to pre-identify candidates having high inhibitory activity, we continued the study with checkerboard experiments. The activities of the compounds with low inhibitory activity in the single-dose experiment were confirmed by this checkerboard analysis. Among those compounds with high inhibitory activity, **2700.001**, **2700.003**, and **2700.004**, only the lattermost did not show inhibition at high levels in the checkerboard assays. These results indicate that the single-dose experiments can be an excellent pre-selection mechanism to eliminate compounds with low activity. Still, the possibility of obtaining false positives makes confirmation by checkerboard studies an essential component of the analysis. It was of interest that a regression analysis considering the molecular docking values (ΔG values) and the compound dose required for a two-fold potentiation as determined by the checkerboard assays, including all compounds tested in both this and the previous study, indicated a significant correlation between ΔG values and inhibitory activity. These results validate docking analysis as an evidentially supportive and potentially predictive tool for further modification that may result in more potent inhibitors.

In conclusion, the results of the structure activity relationship studies indicate that those compounds with the R1-R4 substitutions present in **2700.001** but have modifications at R5 (except for 2-phenylbutyl) are candidates for further modifications at the R3 position. Although no other compound with higher inhibitory activity than the original **2700.001** has been identified, the structure-activity relationship studies have enhanced our understanding of the characteristics and effects of substituting one or more R positions. Moreover, the knowledge acquired directs future analysis to specifically focus on modifying R3, R4, and R5 positions. Overall, the progress achieved in the identification of inhibitors of acetylation mediated by AAC(6′)-Ib by mixture-based combinatorial libraries and application of the positional scanning methodology followed by structure-activity relationship studies with the support of computational molecular modeling, together with excellent work by others to find inhibitors of resistance to a variety of antibiotics [46–48] validates the general strategy as a means to counter antibiotic resistance.

## Declarations

### Funding

*T*his work was supported by Public Health Service grants R15AI047115 (MET) from the National Institute of Allergy and Infectious Diseases, National Institutes of Health and SC3GM125556 (MSR) from the National Institute of General Medicine, National Institutes of Health.

### Competing Interests

None

### Ethical Approval

Not required

## Notes

### Competing Interest Statement

The authors have declared no competing interest.

