## Supplemental file for "Structure-activity relationship of pyrrolidine pentamine derivatives as inhibitors of the aminoglycoside 6′-*N*-acetyltransferase type Ib"

**[2]Center for Translational Science, Florida International University, Port St. Lucie, FL 34987, USA**

**[3]Department of Physics, Florida International University, Miami, FL 33199, USA**

**[4]Biomolecular Sciences Institute, Florida International University, Miami, FL 33199, USA**

**[5]Department of Mathematics, Nova Southeastern University, Fort Lauderdale, FL 33314, USA**

**[6]Medicinal Chemistry, University of Minnesota, Minneapolis, MN 55455, USA**

Figure S1
*Synthetic scheme*

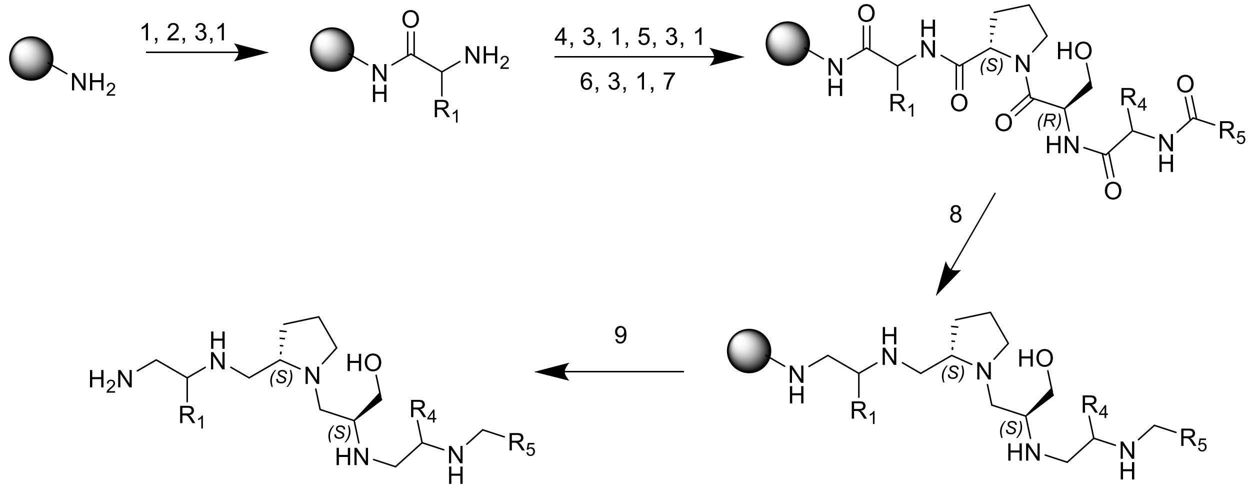


Figure S1. *Synthetic method*. Starting from mBHA resin. 1) 5% DIEA/DCM. 2) 6-fold excess Boc-Amino acid (R1), DIC, and HOBt in 0.1M DMF 2hr. 3) 55% TFA/DCM. 4) 6-fold excess Boc-L-Pro-OH, DIC, and HOBt in 0.1M DMF 2hr. 5) 6-fold excess Boc-D-Ser-OH, DIC, and HOBt in 0.1M DMF 2hr. 6) 6-fold excess Boc-Amino acid (R4), DIC, and HOBt in 0.1M DMF 2hr. 7) 10-fold excess Carboxylic acid (R5), DIC, and HOBt in 0.1M DMF 2hr. 8) 40-fold excess of 0.1M $BH_3$/THF 60°C 72hr followed by Piperidine 60°C 24hr. 9) HF 0°C 7hr.

Figure S2

*Compound synthesis, purification, and characterization.*

All reagents were commercially available and used without further purification. The final compounds were purified using preparative HPLC with a dual pump Shimadzu LC-20AB system equipped with a Luna C18 preparative column (21.5 x 150 mm, 5 micron) at $\lambda$ = 214 nm, with a mobile phase of (A) H2O (+0.1% trifluoroacetic acid)/(B) acetonitrile (ACN) (+0.1% trifluoroacetic acid) at a flow rate of 15 mL/min; gradients varied by compound based on hydrophobicity. The purities of synthesized compounds were confirmed to be greater than 90% by liquid chromatography and mass spectrometry on a Shimadzu LCMS-2020 instrument with ESI Mass Spec and SPD-20A Liquid Chromatograph with a mobile phase of (A) H2O (+0.1% formic acid)/(B) ACN (+0.1% formic acid) (5-95% over 6 min with a 4 min rinse).

Compound **2700-001 (2S)-3-((2S)-2-(((2-amino-1-phenylethyl)amino)methyl)pyrrolidin-1-yl)-2-((2-phenyl-2-((3-phenylbutyl)amino)ethyl)amino)propan-1-ol** Using General Scheme (scheme 1) for the synthesis of pentamines, compound 2700.001 was synthesized using the following reagents: (100mg) MBHA resin starting material, Boc-L-Phenylglycine-OH ($R_1$), Boc-L-Phenylglycine-OH ($R_4$), and 3-Phenylbutyric Acid ($R_5$). The final crude product was purified using HPLC as described above, with a gradient of (B) 0/2, 2/2, 5/5, 40/20, 43/98, 45/98. **Isolated Mass** 4.8mg, **m/z** calculated $C_{34}H_{49}N_5O$ $[M+H]^+$ 543.8, found 544.45 (MS ESI) **Purity** LC-MS: 91.20% (214 nm, peak area).

Compound **2700-002 (2S)-3-((S)-2-((((S)-1-amino-3-phenylpropan-2-yl)amino)methyl)pyrrolidin-1-yl)-2-((2-phenyl-2-((3-phenylbutyl)amino)ethyl)amino)propan-1-ol.** Using General Scheme (scheme 1) for the synthesis of pentamines, compound 2700.002 was synthesized using the following reagents: (100mg) MBHA resin starting material, Boc-L-Phenylalanine-OH ($R_1$), Boc-L-Phenylglycine-OH ($R_4$), and 3-Phenylbutyric Acid ($R_5$). The final crude product was purified using HPLC as described above, with a gradient of (B) 0/2, 2/2, 5/5, 40/20, 43/98, 45/98. **Isolated Mass** 8.6mg, **m/z** calculated $C_{35}H_{51}N_5O$ $[M+H]^+$ 557.83, found 558.45 (MS ESI) **Purity** LC-MS: 90.10% (214 nm, peak area).

Compound **2700-003 (2S)-3-((2S)-2-(((2-amino-1-phenylethyl)amino)methyl)pyrrolidin-1-yl)-2-((2-phenyl-2-((4-phenylbutyl)amino)ethyl)amino)propan-1-ol.** Using General Scheme (scheme 1) for the synthesis of pentamines, compound 2700.003 was synthesized using the following reagents: (100mg) MBHA resin starting material, Boc-L-Phenylglycine-OH ($R_1$), Boc-L-Phenylglycine-OH ($R_4$), and 4-Phenylbutyric Acid ($R_5$). The final crude product was purified using HPLC as described above, with a gradient of (B) 0/2, 2/2, 5/5, 40/20, 43/98, 45/98. **Isolated Mass** 5.1mg, **m/z** calculated $C_{34}H_{49}N_5$ $[M+H]^+$ 543.8, found 544.45 (MS ESI) **Purity** LC-MS: 90.01% (214 nm, peak area).

Figure S2 continued

Compound **2700-004 (2S)-3-((2S)-2-(((2-amino-1-phenylethyl)amino)methyl)pyrrolidin-1-yl)-2-(((S)-3-phenyl-2-((4-phenylbutyl)amino)propyl)amino)propan-1-ol.** Using General Scheme (scheme 1) for the synthesis of pentamines, compound 2700.004 was synthesized using the following reagents: (100mg) MBHA resin starting material, Boc-L-Phenylglycine-OH ($R_1$), Boc-L-Phenylalanine-OH ($R_4$), and 4-Phenylbutyric Acid ($R_5$). The final crude product was purified using HPLC as described above, with a gradient of (B) 0/2, 2/2, 5/5, 40/20, 43/98, 45/98. **Isolated Mass** 5.5mg, **m/z** calculated $C_{35}H_{51}N_5O$ $[M+H]^+$ 557.83, found 558.4 (MS ESI) **Purity** LC-MS: 92.19% (214 nm, peak area).

Compound **2700-005 (2S)-3-((2S)-2-(((2-amino-1-phenylethyl)amino)methyl)pyrrolidin-1-yl)-2-(((2S)-3-phenyl-2-((3-phenylbutyl)amino)propyl)amino)propan-1-ol.** Using General Scheme (scheme 1) for the synthesis of pentamines, compound 2700.005 was synthesized using the following reagents: (100mg) MBHA resin starting material, Boc-L-Phenylglycine-OH ($R_1$), Boc-L-Phenylalanine-OH ($R_4$), and 3-Phenylbutyric Acid ($R_5$). The final crude product was purified using HPLC as described above, with a gradient of (B) 0/2, 2/2, 5/5, 40/20, 43/98, 45/98. **Isolated Mass** 5.6mg, **m/z** calculated $C_{35}H_{51}N_5O$ $[M+H]^+$ 557.83, found 558.4 (MS ESI) **Purity** LC-MS: 90.59% (214 nm, peak area).

Compound **2700-006 (2S)-3-((S)-2-((((S)-1-amino-3-(pyridin-3-yl)propan-2-yl)amino)methyl)pyrrolidin-1-yl)-2-((2-phenyl-2-((3-phenylbutyl)amino)ethyl)amino)propan-1-ol.** Using General Scheme (scheme 1) for the synthesis of pentamines, compound 2700.006 was synthesized using the following reagents: (100mg) MBHA resin starting material, Boc-L-Alanine(3-pyridyl)-OH ($R_1$), Boc-L-Phenylglycine-OH ($R_4$), and 3-Phenylbutyric Acid ($R_5$). The final crude product was purified using HPLC as described above, with a gradient of (B) 0/2, 2/2, 5/5, 40/20, 43/98, 45/98. **Isolated Mass** 5.6mg, **m/z** calculated $C_{34}H_{50}N_6O$ $[M+H]^+$ 558.82, found 559.45 (MS ESI) **Purity** LC-MS: 92.18% (214 nm, peak area).

Compound **2700-007 (2S)-3-((S)-2-((((S)-1-amino-3-(pyridin-3-yl)propan-2-yl)amino)methyl)pyrrolidin-1-yl)-2-((2-phenyl-2-((4-phenylbutyl)amino)ethyl)amino)propan-1-ol.** Using General Scheme (scheme 1) for the synthesis of pentamines, compound 2700.007 was synthesized using the following reagents: (100mg) MBHA resin starting material, Boc-L-Alanine(3-pyridyl)-OH ($R_1$), Boc-L-Phenylglycine-OH ($R_4$), and 4-Phenylbutyric Acid ($R_5$). The final crude product was purified using HPLC as described above, with a gradient of (B) 0/2, 2/2, 5/5, 40/20, 43/98, 45/98. **Isolated Mass** 6.4mg, **m/z** calculated $C_{34}H_{50}N_6O$ $[M+H]^+$ 558.82, found 559.4 (MS ESI) **Purity** LC-MS: 84.46% (214 nm, peak area).

Figure S2 continued

Compound **2700-008** **(S)-3-((S)-2-((((S)-1-amino-3-(pyridin-3-yl)propan-2-yl)amino)methyl)pyrrolidine-1-yl)-2-(((S)-3-phenyl-2-((4-phenylbutyl)amino)propyl)amino)propan-1-ol.** Using General Scheme (scheme 1) for the synthesis of pentamines, compound 2700.008 was synthesized using the following reagents: (100mg) MBHA resin starting material, Boc-L-Alanine(3-pyridyl)-OH ($R_1$), Boc-L-Phenylalanine-OH ($R_4$), and 4-Phenylbutyric Acid ($R_5$). The final crude product was purified using HPLC as described above, with a gradient of (B) 0/2, 2/2, 5/5, 40/20, 43/98, 45/98. **Isolated Mass 6.4**mg, **m/z** calculated $C_{35}H_{52}N_6O$ $[M+H]^+$ 572.84, found 573.45 (MS ESI) **Purity** LC-MS: 96.22% (214 nm, peak area).

Compound **2700-009** **(2S)-3-((S)-2-((((S)-1-amino-3-phenylpropan-2-yl)amino)methyl)pyrrolidin-1-yl)-2-((2-phenyl-2-((4-phenylbutyl)amino)ethyl)amino)propan-1-ol.** Using General Scheme (scheme 1) for the synthesis of pentamines, compound 2700.009 was synthesized using the following reagents: (100mg) MBHA resin starting material, Boc-L-Phenylalanine-OH ($R_1$), Boc-L-Phenylglycine-OH ($R_4$), and 4-Phenylbutyric Acid ($R_5$). The final crude product was purified using HPLC as described above, with a gradient of (B) 0/2, 2/2, 5/5, 40/20, 43/98, 45/98. **Isolated Mass** 6.1mg, **m/z** calculated $C_{35}H_{51}N_5O$ $[M+H]^+$ 557.83, found 558.4 (MS ESI) **Purity** LC-MS: 92.26% (214 nm, peak area).

Compound **2700-010** **(S)-3-((S)-2-((((S)-1-amino-3-phenylpropan-2-yl)amino)methyl)pyrrolidin-1-yl)-2-(((S)-3-phenyl-2-((4-phenylbutyl)amino)propyl)amino)propan-1-ol.** Using General Scheme (scheme 1) for the synthesis of pentamines, compound 2700.010 was synthesized using the following reagents: (100mg) MBHA resin starting material, Boc-L-Phenylalanine-OH ($R_1$), Boc-L-Phenylalanine-OH ($R_4$), and 4-Phenylbutyric Acid ($R_5$). The final crude product was purified using HPLC as described above, with a gradient of (B) 0/2, 2/2, 5/5, 40/20, 43/98, 45/98. **Isolated Mass** 6.1mg, **m/z** calculated $C_{36}H_{53}N_5O$ $[M+H]^+$ 571.85, found 572.5 (MS ESI) **Purity** LC-MS: 93.97% (214 nm, peak area).

Compound **2700-011** **4-((2S)-3-amino-2-((((2S)-1-((2S)-3-hydroxy-2-((2-phenyl-2-((3-phenylbutyl)amino)ethyl)amino)propyl)pyrrolidin-2-yl)methyl)amino)propyl)phenol.** Using General Scheme (scheme 1) for the synthesis of pentamines, compound 2700.011 was synthesized using the following reagents: (100mg) MBHA resin starting material, Boc-L-Tyrosine-OH ($R_1$), Boc-L-Phenylglycine-OH ($R_4$), and 3-Phenylbutyric Acid ($R_5$). The final crude product was purified using HPLC as described above, with a gradient of (B) 0/2, 2/2, 5/5, 40/20, 43/98, 45/98. **Isolated Mass** 5.6mg, **m/z** calculated $C_{35}H_{51}N_5O_2$ $[M+H]^+$ 573.83, found 574.45 (MS ESI) **Purity** LC-MS: 92.79% (214 nm, peak area).

Figure S2 continued

Compound **2700-012 4-((2S)-3-amino-2-((((2S)-1-((2S)-3-hydroxy-2-((2-phenyl-2-((4-phenylbutyl)amino)ethyl)amino)propyl)pyrrolidin-2-yl)methyl)amino)propyl)phenol.** Using General Scheme (scheme 1) for the synthesis of pentamines, compound 2700.012 was synthesized using the following reagents: (100mg) MBHA resin starting material, Boc-L-Tyrosine-OH ($R_1$), Boc-L-Phenylglycine-OH ($R_4$), and 4-Phenylbutyric Acid ($R_5$). The final crude product was purified using HPLC as described above, with a gradient of (B) 0/2, 2/2, 5/5, 40/20, 43/98, 45/98. **Isolated Mass** 4.7mg, **m/z** calculated $C_{35}H_{51}N_5O_2$ $[M+H]^+$ 573.83, found 574.45 (MS ESI) **Purity** LC-MS: 92.94% (214 nm, peak area).

Compound **2700-013 4-((S)-3-amino-2-((((S)-1-((S)-3-hydroxy-2-(((S)-3-phenyl-2-((4-phenylbutyl)amino)propyl)amino)propyl)pyrrolidin-2-yl)methyl)amino)propyl)phenol.** Using General Scheme (scheme 1) for the synthesis of pentamines, compound 2700.013 was synthesized using the following reagents: (100mg) MBHA resin starting material, Boc-L-Tyrosine-OH ($R_1$), Boc-L-Phenylalanine-OH ($R_4$), and 4-Phenylbutyric Acid ($R_5$). The final crude product was purified using HPLC as described above, with a gradient of (B) 0/2, 2/2, 5/5, 40/20, 43/98, 45/98. **Isolated Mass** 6.0mg, **m/z** calculated $C_{36}H_{53}N_5O_2$ $[M+H]^+$ 587.85, found 588.25 (MS ESI) **Purity** LC-MS: 92.59% (214 nm, peak area).

Compound **2700-014 (2S)-3-((S)-2-((((R)-1-aminohexan-2-yl)amino)methyl)pyrrolidin-1-yl)-2-((2-phenyl-2-((4-phenylbutyl)amino)ethyl)amino)propan-1-ol.** Using General Scheme (scheme 1) for the synthesis of pentamines, compound 2700.014 was synthesized using the following reagents: (100mg) MBHA resin starting material, Boc-D-Norleucine-OH ($R_1$), Boc-L-Phenylglycine-OH ($R_4$), and 4-Phenylbutyric Acid ($R_5$). The final crude product was purified using HPLC as described above, with a gradient of (B) 0/2, 2/2, 5/5, 40/20, 43/98, 45/98. **Isolated Mass** 5.1mg, **m/z** calculated $C_{32}H_{53}N_5O$ $[M+H]^+$ 523.81, found 524.25 (MS ESI) **Purity** LC-MS: 76.03% (214 nm, peak area).

Compound **2700-015 (S)-3-((S)-2-((((R)-1-aminohexan-2-yl)amino)methyl)pyrrolidin-1-yl)-2-(((S)-3-phenyl-2-((4-phenylbutyl)amino)propyl)amino)propan-1-ol.** Using General Scheme (scheme 1) for the synthesis of pentamines, compound 2700.015 was synthesized using the following reagents: (100mg) MBHA resin starting material, Boc-D-Norleucine-OH ($R_1$), and Boc-L-Phenylalanine-OH ($R_4$), and 4-Phenylbutyric Acid ($R_5$). The final crude product was purified using HPLC as described above, with a gradient of (B) 0/2, 2/2, 5/5, 40/20, 43/98, 45/98. **Isolated Mass** 5.4mg, **m/z** calculated $C_{33}H_{55}N_5O$ $[M+H]^+$ 537.84, found 538.25 (MS ESI) **Purity** LC-MS: 82.97% (214 nm, peak area).

Figure S3

*Liquid chromatography-mass spectrometry analysis*

**2700-001**



#### <Sample Information>

| | | | |
|---|---|---|---|
| Sample Name | : 2700 | | |
| Sample ID | : 2700-A1 peak 1 | | |
| Data Filename | : 2700-A1 peak 1 080421.lcd | | |
| Method Filename | : 6min 5-95, 1min clean(254)_(214) Col_1_LCMS3.lcm | | |
| Batch Filename | : 080421.lcb | | |
| Vial # | : 1-1 | Sample Type | : Unknown |
| Injection Volume | : 10 uL | | |
| Date Acquired | : 8/4/2021 4:12:21 PM | Acquired by | : System Administrator |
| Date Processed | : 9/1/2021 3:08:34 PM | Processed by | : System Administrator |

#### <Chromatogram>

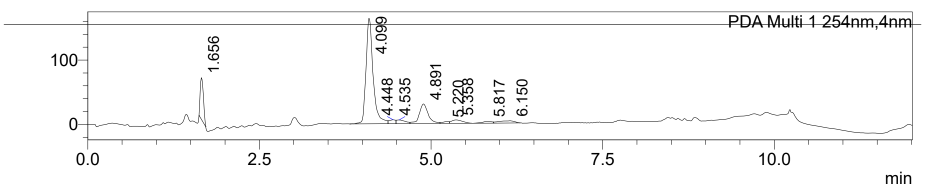


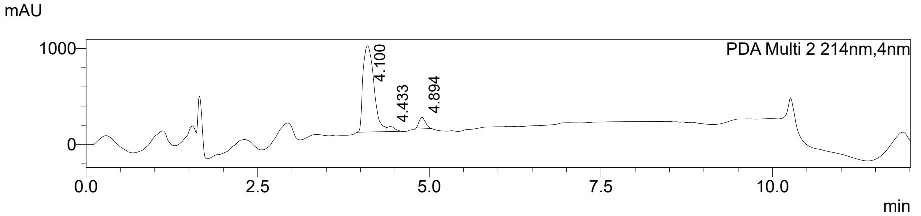


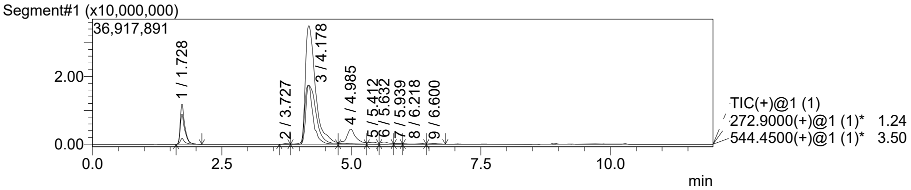


MS Spectrum

Peak#:1 R.Time:1.728(Scan#:161)
MassPeaks:72
Spectrum Mode:Averaged 1.723-1.744(160-162)
BG Mode:Calc Segment 1 - Event 1

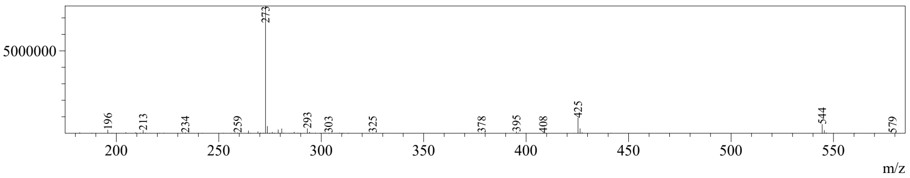

Figure S3 continued

**2700-002**



### <Sample Information>

| | | | |
|---|---|---|---|
| Sample Name | : 2700 | | |
| Sample ID | : 2700-A2 peak 1 | | |
| Data Filename | : 2700-A2 peak 1 080421.lcd | | |
| Method Filename | : 6min 5-95, 1min clean(254)_(214) Col_1_LCMS3.lcm | | |
| Batch Filename | : 080421.lcb | | |
| Vial # | : 1-5 | Sample Type | : Unknown |
| Injection Volume | : 10 uL | | |
| Date Acquired | : 8/4/2021 5:02:19 PM | Acquired by | : System Administrator |
| Date Processed | : 4/16/2024 2:37:46 PM | Processed by | : System Administrator |

### <Chromatogram>

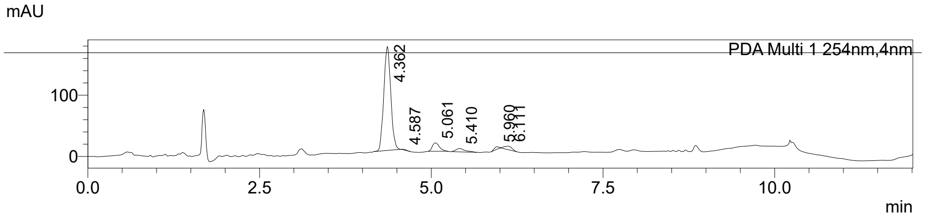


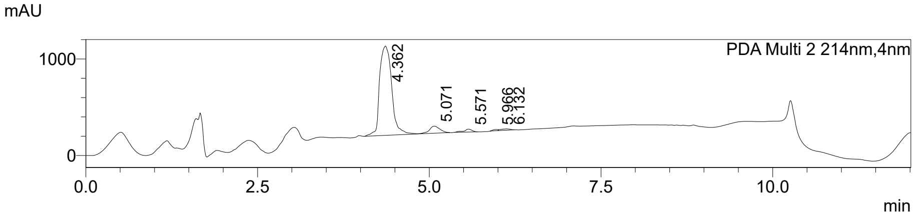


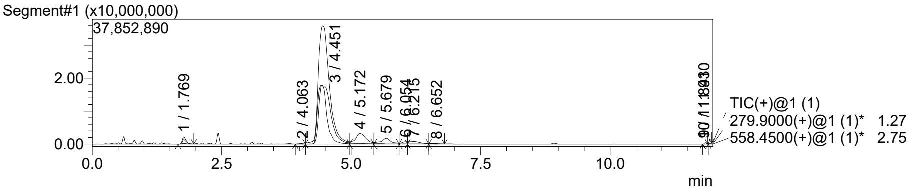


MS Spectrum

Peak#:1  R.Time:1.769(Scan#:164)
MassPeaks:54
Spectrum Mode:Averaged 1.755-1.777(163-165)
BG Mode:Calc  Segment 1 - Event 1

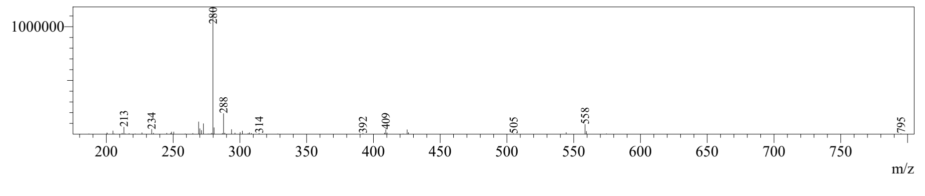

Figure S3 continued

**2700-003**



**<Sample Information>**

| | | | |
|---|---|---|---|
| Sample Name | : 2700 | | |
| Sample ID | : 2700-A3 peak 1 | | |
| Data Filename | : 2700-A3 peak 1 080421.lcd | | |
| Method Filename | : 6min 5-95, 1min clean(254)_(214) Col_1_LCMS3.lcm | | |
| Batch Filename | : 080421.lcb | | |
| Vial # | : 1-7 | Sample Type | : Unknown |
| Injection Volume | : 10 uL | | |
| Date Acquired | : 8/4/2021 5:27:19 PM | Acquired by | : System Administrator |
| Date Processed | : 4/16/2024 2:46:33 PM | Processed by | : System Administrator |

**<Chromatogram>**

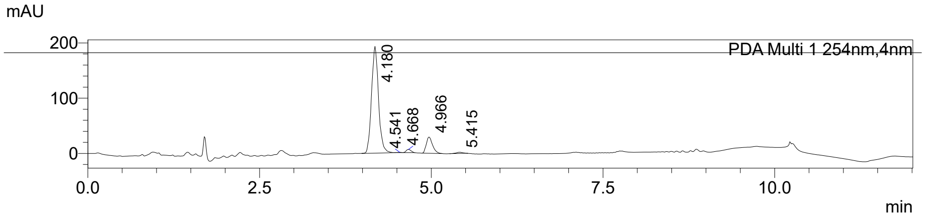


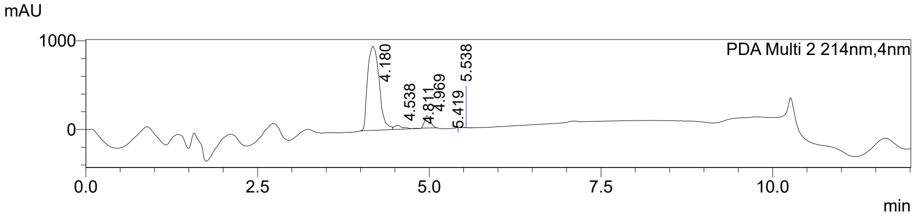


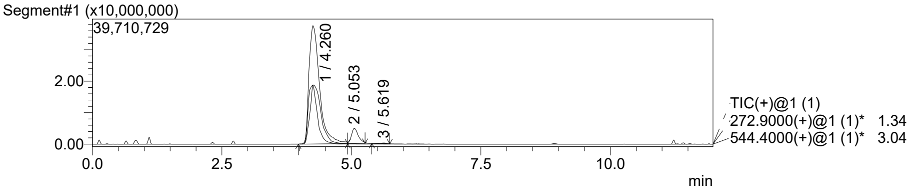


MS Spectrum

Peak#:1 R.Time:4.260(Scan#:394)
MassPeaks:71
Spectrum Mode:Averaged 4.247-4.268(393-395)
BG Mode:Calc Segment 1 - Event 1

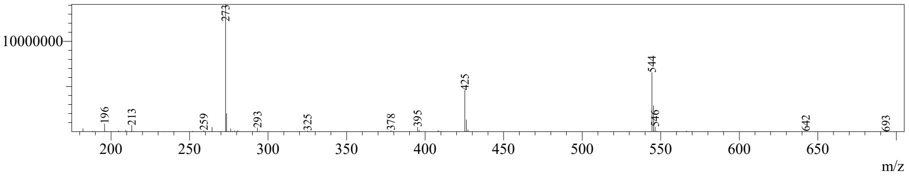

Figure S3 continued

**2700-004**



**<Sample Information>**

| | | | |
|---|---|---|---|
| Sample Name | : 2700 | | |
| Sample ID | : 2700-A4 peak 1 | | |
| Data Filename | : 2700-A4 peak 1 080421.lcd | | |
| Method Filename | : 6min 5-95, 1min clean(254)_(214) Col_1_LCMS3.lcm | | |
| Batch Filename | : 080421.lcb | | |
| Vial # | : 1-10 | Sample Type | : Unknown |
| Injection Volume | : 10 uL | | |
| Date Acquired | : 8/4/2021 6:04:50 PM | Acquired by | : System Administrator |
| Date Processed | : 9/1/2021 3:35:47 PM | Processed by | : System Administrator |

**<Chromatogram>**

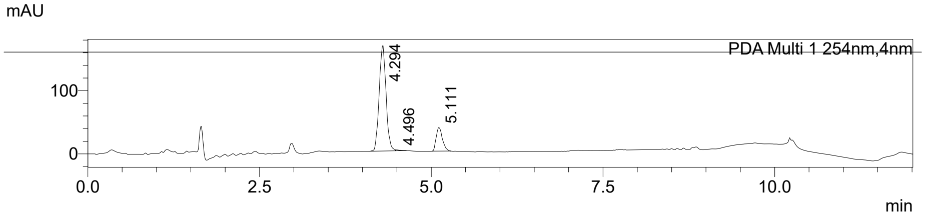


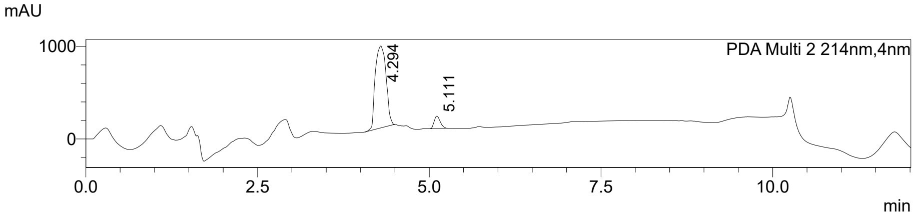


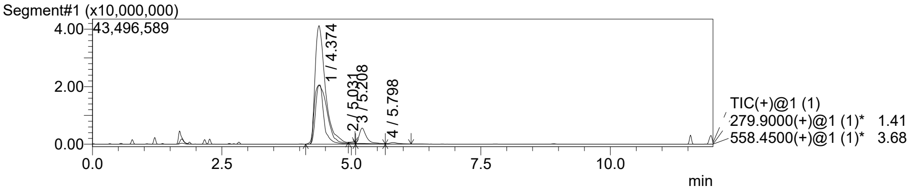


MS Spectrum

Peak#:1 R.Time:4.374(Scan#:405)
MassPeaks:77
Spectrum Mode:Averaged 4.366-4.388(404-406)
BG Mode:Calc Segment 1 - Event 1

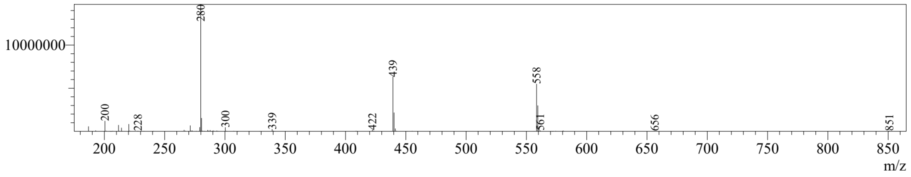

Figure S3 continued

**2700-005**



### <Sample Information>

| | | | |
|---|---|---|---|
| Sample Name | : 2700 | | |
| Sample ID | : 2700-A5 peak 1 | | |
| Data Filename | : 2700-A5 peak 1 080421.lcd | | |
| Method Filename | : 6min 5-95, 1min clean(254)_(214) Col_1_LCMS3.lcm | | |
| Batch Filename | : 080421.lcb | | |
| Vial # | : 1-13 | Sample Type | : Unknown |
| Injection Volume | : 10 uL | | |
| Date Acquired | : 8/4/2021 6:54:32 PM | Acquired by | : System Administrator |
| Date Processed | : 9/1/2021 3:37:37 PM | Processed by | : System Administrator |

### <Chromatogram>

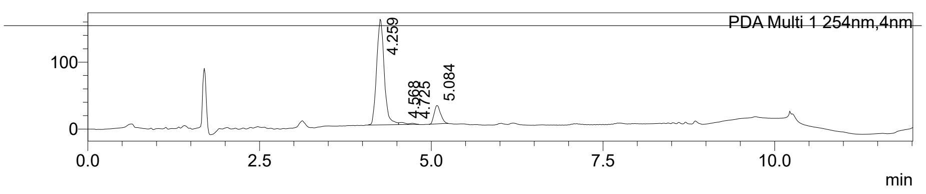


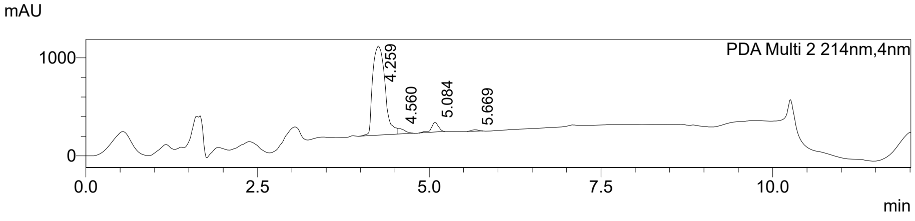


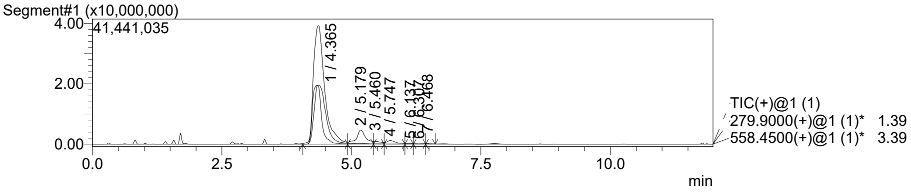


MS Spectrum

Peak#:1 R.Time:4.365(Scan#:404)
MassPeaks:74
Spectrum Mode:Averaged 4.355-4.377(403-405)
BG Mode:Calc Segment 1 - Event 1

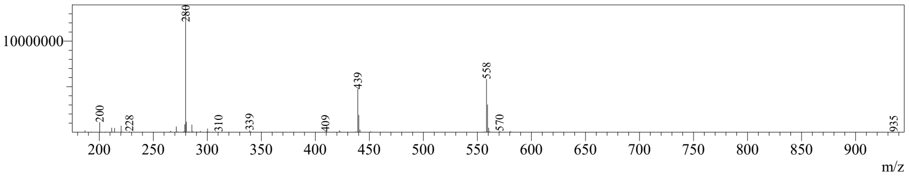

Figure S3 continued
**2700-006**



### <Sample Information>

| | | | |
|---|---|---|---|
| Sample Name | : 2700 | | |
| Sample ID | : 2700-A6 peak 1 | | |
| Data Filename | : 2700-A6 peak 1 080521.lcd | | |
| Method Filename | : 6min 5-95, 1min clean(254)_(214) Col_1_LCMS3.lcm | | |
| Batch Filename | : 080521.lcb | | |
| Vial # | : 1-16 | Sample Type | : Unknown |
| Injection Volume | : 5 uL | | |
| Date Acquired | : 8/5/2021 4:44:08 PM | Acquired by | : System Administrator |
| Date Processed | : 9/1/2021 3:39:08 PM | Processed by | : System Administrator |

### <Chromatogram>

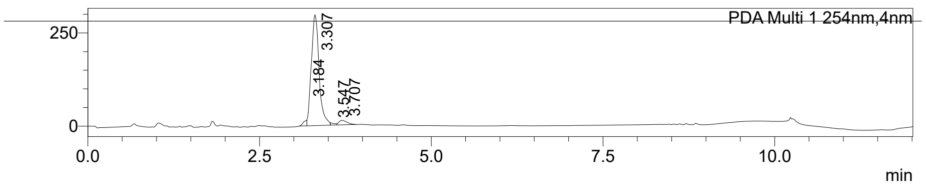


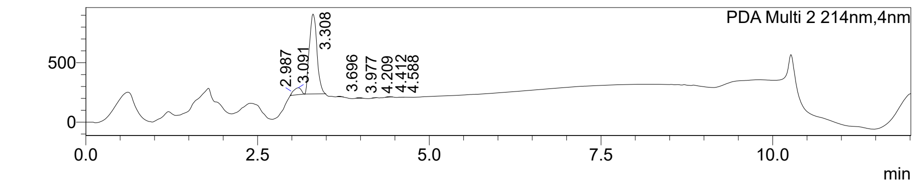


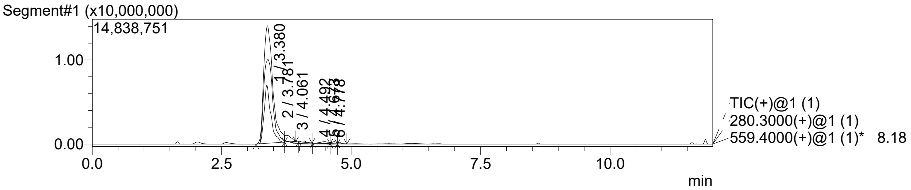


MS Spectrum

Peak#:1 R.Time:3.380(Scan#:313)
MassPeaks:69
Spectrum Mode:Averaged 3.369-3.391(312-314)
BG Mode:Calc Segment 1 - Event 1

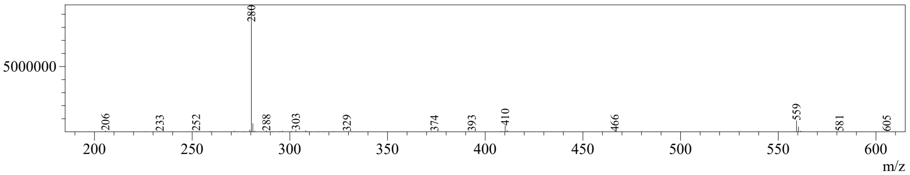

Figure S3 continued

**2700-007**



**<Sample Information>**

| | | | |
|---|---|---|---|
| Sample Name | : 2700 | | |
| Sample ID | : 2700-A7 peak 1 | | |
| Data Filename | : 2700-A7 peak 1 080421b.lcd | | |
| Method Filename | : 6min 5-95, 1min clean(254)_(214) Col_1_LCMS3.lcm | | |
| Batch Filename | : 080421.lcb | | |
| Vial # | : 1-19 | Sample Type | : Unknown |
| Injection Volume | : 10 uL | | |
| Date Acquired | : 8/4/2021 8:09:31 PM | Acquired by | : System Administrator |
| Date Processed | : 9/1/2021 4:17:17 PM | Processed by | : System Administrator |

**<Chromatogram>**

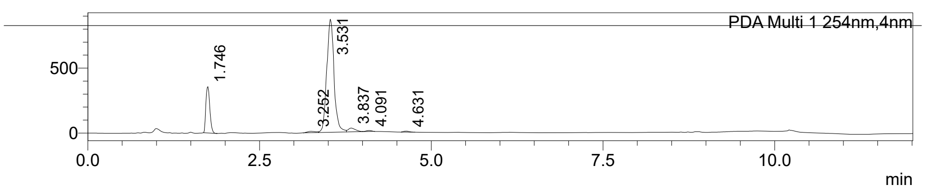


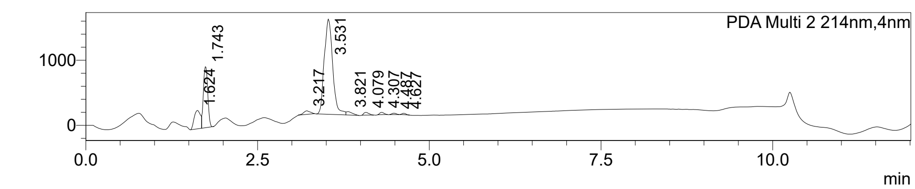


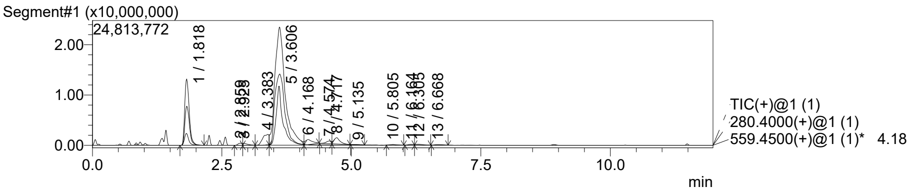


MS Spectrum

Peak#:1 R.Time:1.818(Scan#:169)
MassPeaks:115
Spectrum Mode:Averaged 1.809-1.831(168-170)
BG Mode:Calc Segment 1 - Event 1

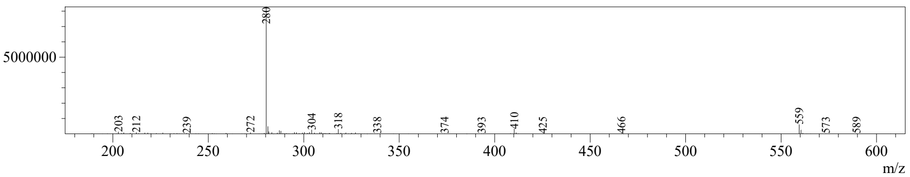

Figure S3 continued
**2700-008**



**<Sample Information>**

Sample Name : 2700
Sample ID : 2700-A8 peak 1
Data Filename : 2700-A8 peak 1 080421.lcd
Method Filename : 6min 5-95, 1min clean(254)_(214) Col_1_LCMS3.lcm
Batch Filename : 080421.lcb
Vial # : 1-21 Sample Type : Unknown
Injection Volume : 10 uL
Date Acquired : 8/4/2021 8:46:39 PM Acquired by : System Administrator
Date Processed : 9/1/2021 4:24:48 PM Processed by : System Administrator

**<Chromatogram>**

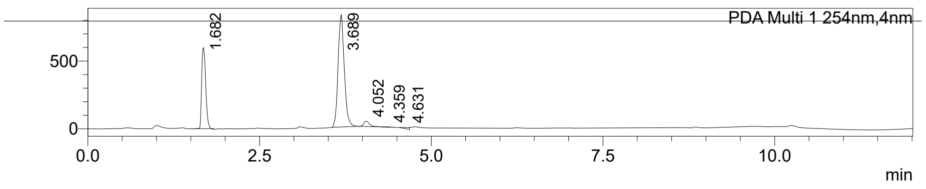


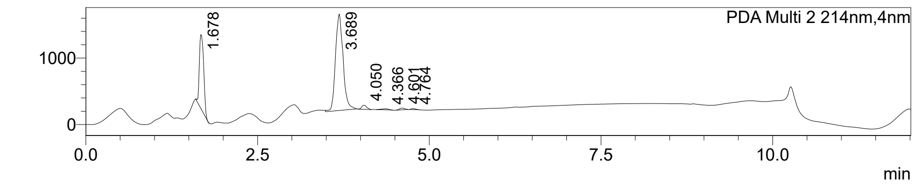


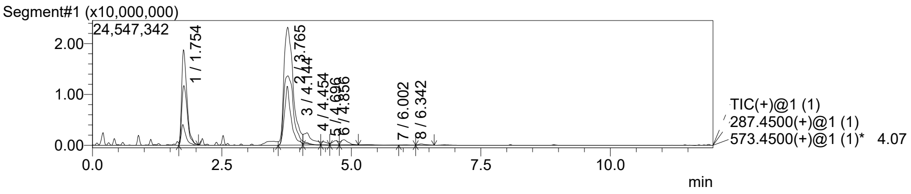


MS Spectrum
Peak#:1 R.Time:1.754(Scan#:163)
MassPeaks:127
Spectrum Mode:Averaged 1.744-1.766(162-164)
BG Mode:Calc Segment 1 - Event 1

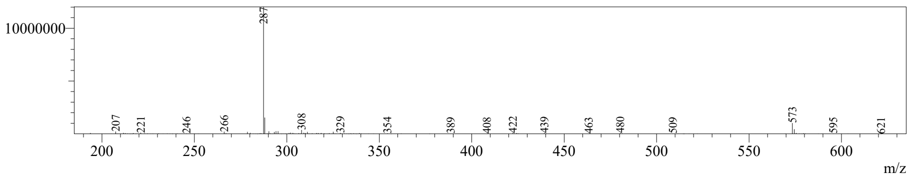

Figure S3 continued

**2700-009**



**<Sample Information>**

| | | | |
|---|---|---|---|
| Sample Name | : 2700 | | |
| Sample ID | : 2700-A9 peak 1 | | |
| Data Filename | : 2700-A9 peak 1 080421.lcd | | |
| Method Filename | : 6min 5-95, 1min clean(254)_(214) Col_1_LCMS3.lcm | | |
| Batch Filename | : 080421.lcb | | |
| Vial # | : 1-22 | Sample Type | : Unknown |
| Injection Volume | : 10 uL | | |
| Date Acquired | : 8/4/2021 8:59:08 PM | Acquired by | : System Administrator |
| Date Processed | : 9/1/2021 4:54:05 PM | Processed by | : System Administrator |

**<Chromatogram>**

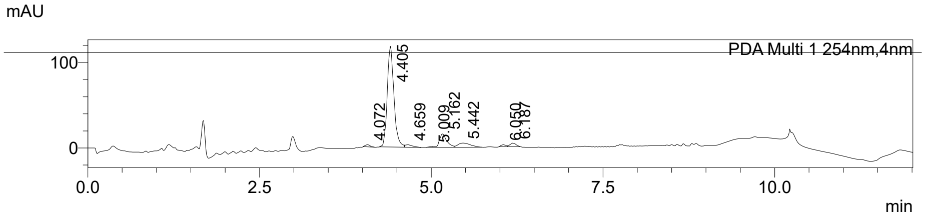


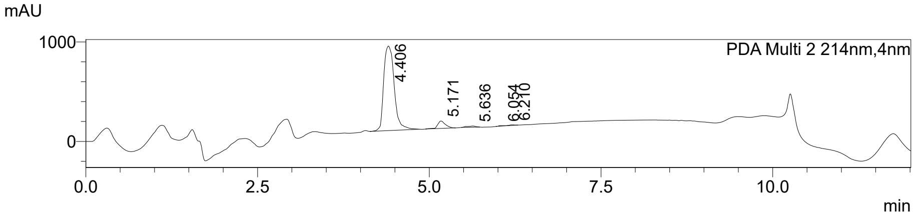


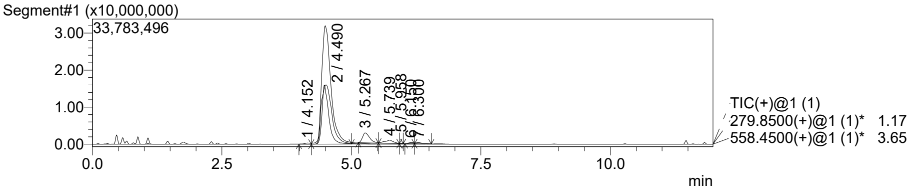


MS Spectrum

Peak#:1 R.Time:4.152(Scan#:384)
MassPeaks:8
Spectrum Mode:Averaged 4.138-4.160(383-385)
BG Mode:Calc Segment 1 - Event 1

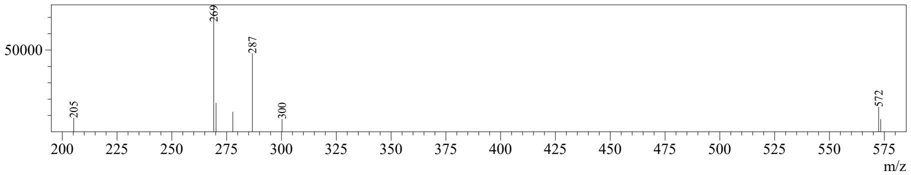

Figure S3 continued

**2700-010**



### <Sample Information>

| | | | |
|---|---|---|---|
| Sample Name | : 2700 | | |
| Sample ID | : 2700-A10 peak 1 | | |
| Data Filename | : 2700-A10 peak 1 080421.lcd | | |
| Method Filename | : 6min 5-95, 1min clean(254)_(214) Col_1_LCMS3.lcm | | |
| Batch Filename | : 080421.lcb | | |
| Vial # | : 1-23 | Sample Type | : Unknown |
| Injection Volume | : 10 uL | | |
| Date Acquired | : 8/4/2021 9:11:37 PM | Acquired by | : System Administrator |
| Date Processed | : 9/1/2021 5:42:10 PM | Processed by | : System Administrator |

### <Chromatogram>

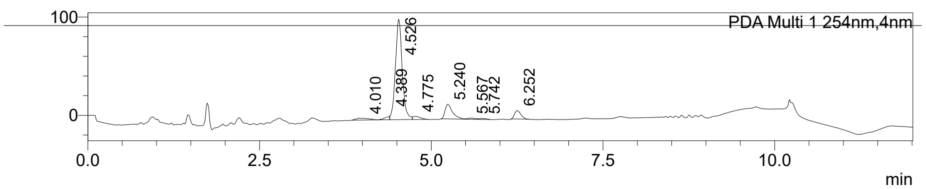


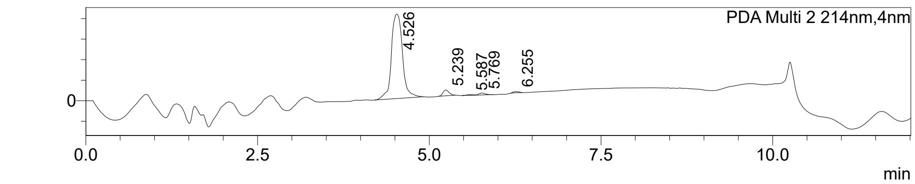


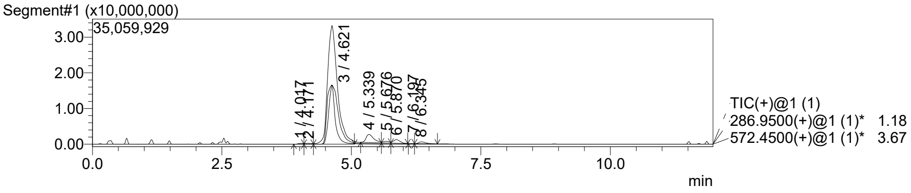


MS Spectrum

Peak#:1 R.Time:4.017(Scan#:372)
MassPeaks:7
Spectrum Mode:Averaged 4.008-4.030(371-373)
BG Mode:Calc Segment 1 - Event 1

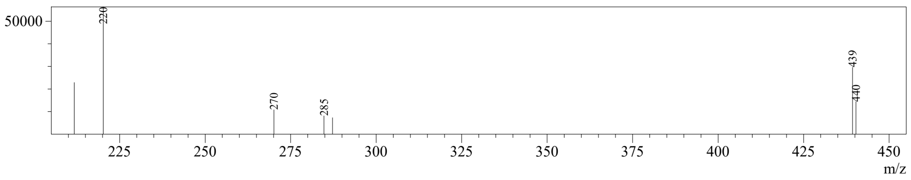

Figure S3 continued

**2700-011**



### <Sample Information>

| | | | |
|---|---|---|---|
| Sample Name | : 2700 | | |
| Sample ID | : 2700-A11 peak 1 | | |
| Data Filename | : 2700-A11 peak 1 080521.lcd | | |
| Method Filename | : 6min 5-95, 1min clean(254)_(214) Col_1_LCMS3.lcm | | |
| Batch Filename | : 080421.lcb | | |
| Vial # | : 1-26 | Sample Type | : Unknown |
| Injection Volume | : 5 uL | | |
| Date Acquired | : 8/5/2021 1:11:03 PM | Acquired by | : System Administrator |
| Date Processed | : 9/7/2021 12:12:59 PM | Processed by | : System Administrator |

### <Chromatogram>

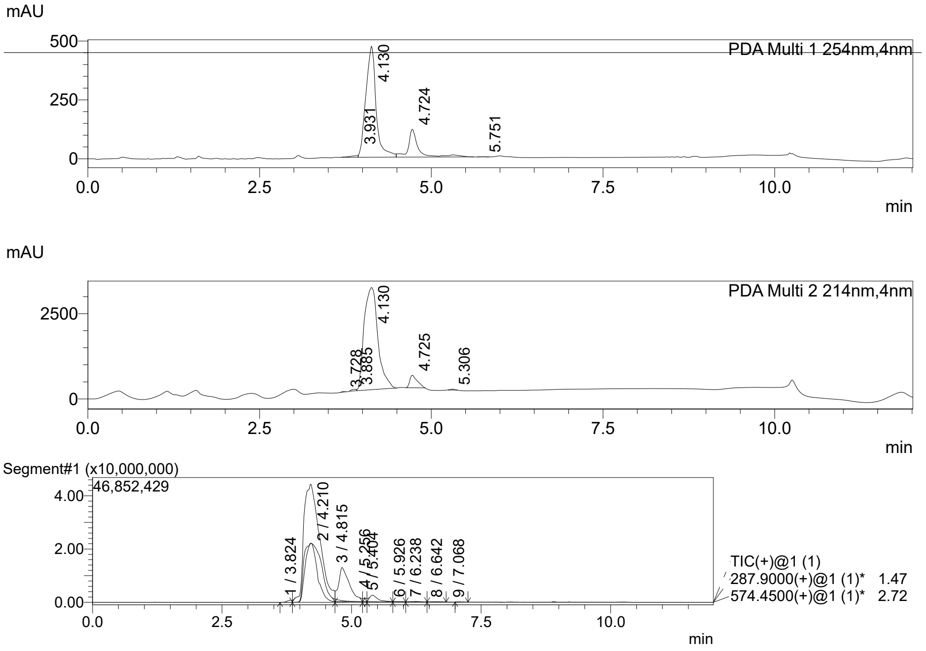


MS Spectrum

Peak#:1 R.Time:3.824(Scan#:354)
MassPeaks:11
Spectrum Mode:Averaged 3.813-3.835(353-355)
BG Mode:Calc Segment 1 - Event 1

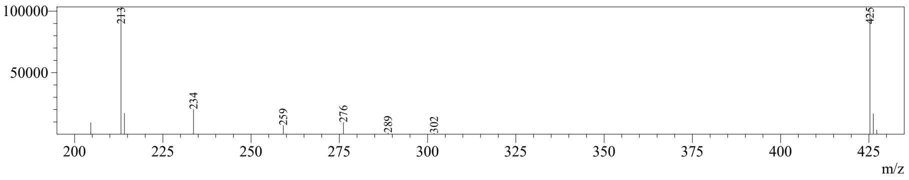

Figure S3 continued

**2700-012**



#### <Sample Information>

| | | | |
|---|---|---|---|
| Sample Name | : 2700 | | |
| Sample ID | : 2700-A12 peak 1 | | |
| Data Filename | : 2700-A12 peak 1 080421.lcd | | |
| Method Filename | : 6min 5-95, 1min clean(254)_(214) Col_1_LCMS3.lcm | | |
| Batch Filename | : 080421.lcb | | |
| Vial # | : 1-29 | Sample Type | : Unknown |
| Injection Volume | : 10 uL | | |
| Date Acquired | : 8/4/2021 10:26:37 PM | Acquired by | : System Administrator |
| Date Processed | : 8/5/2021 3:29:29 PM | Processed by | : System Administrator |

#### <Chromatogram>

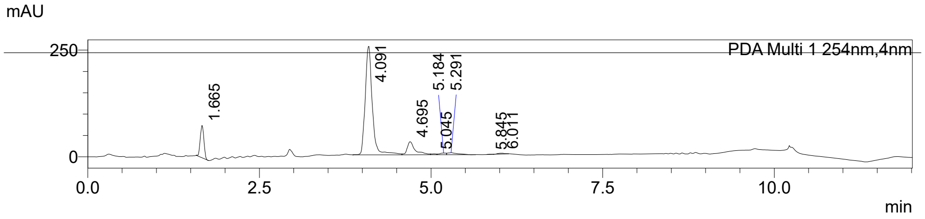


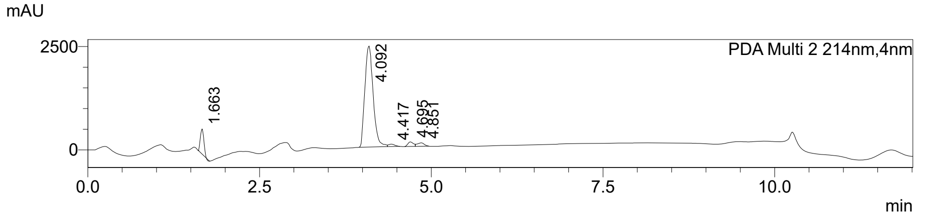


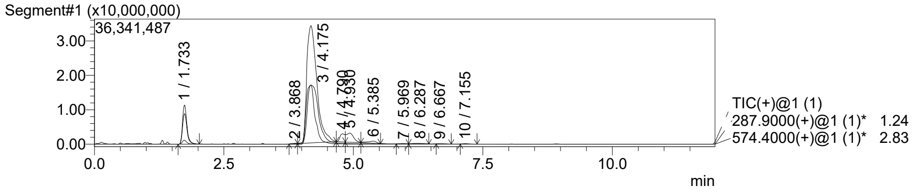


MS Spectrum

Peak#:1 R.Time:1.733(Scan#:161)
MassPeaks:67
Spectrum Mode:Averaged 1.723-1.744(160-162)
BG Mode:Calc Segment 1 - Event 1

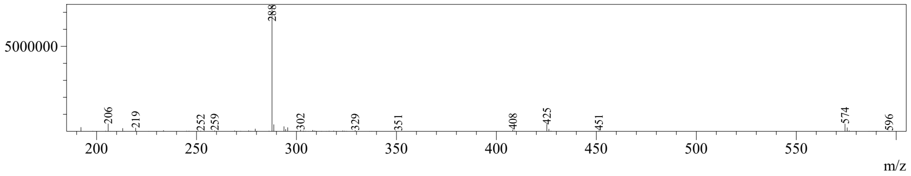

Figure S3 continued

**2700-013**



**<Sample Information>**

| | | | |
|---|---|---|---|
| Sample Name | : 2700 | | |
| Sample ID | : 2700-A13 peak 1 | | |
| Data Filename | : 2700-A13 peak 1 080421.lcd | | |
| Method Filename | : 6min 5-95, 1min clean(254)_(214) Col_1_LCMS3.lcm | | |
| Batch Filename | : 080421.lcb | | |
| Vial # | : 1-30 | Sample Type | : Unknown |
| Injection Volume | : 10 uL | | |
| Date Acquired | : 8/5/2021 10:04:14 AM | Acquired by | : System Administrator |
| Date Processed | : 4/17/2024 2:34:20 PM | Processed by | : System Administrator |

**<Chromatogram>**

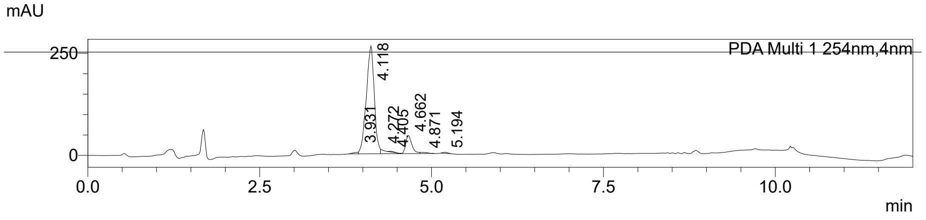


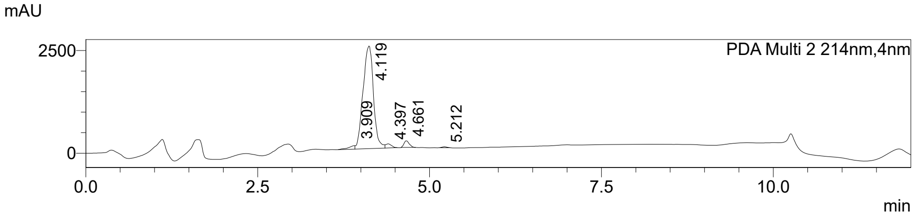


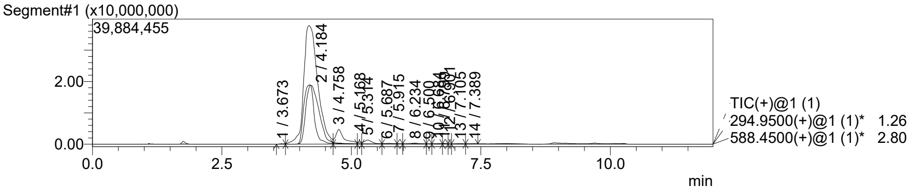


MS Spectrum

Peak#:1 R.Time:3.673(Scan#:340)
MassPeaks:4
Spectrum Mode:Averaged 3.662-3.683(339-341)
BG Mode:Calc Segment 1 - Event 1

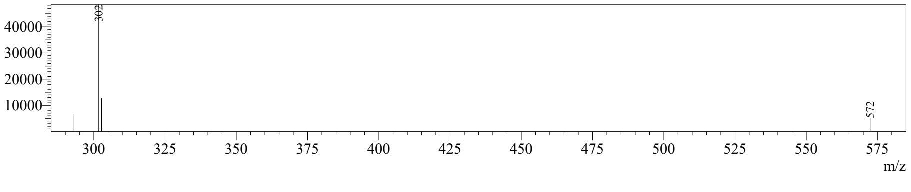

Figure S3 continued
**2700-014**



**<Sample Information>**

| | | | |
|---|---|---|---|
| Sample Name | : 2700 | | |
| Sample ID | : 2700-A14 peak 1 <90 | | |
| Data Filename | : 2700-A14 peak 1 less than 90 080421.lcd | | |
| Method Filename | : 6min 5-95, 1min clean(254)_(214) Col_1_LCMS3.lcm | | |
| Batch Filename | : 080421.lcb | | |
| Vial # | : 1-32 | Sample Type | : Unknown |
| Injection Volume | : 10 uL | | |
| Date Acquired | : 8/5/2021 10:41:25 AM | Acquired by | : System Administrator |
| Date Processed | : 9/20/2021 9:36:13 PM | Processed by | : System Administrator |

**<Chromatogram>**

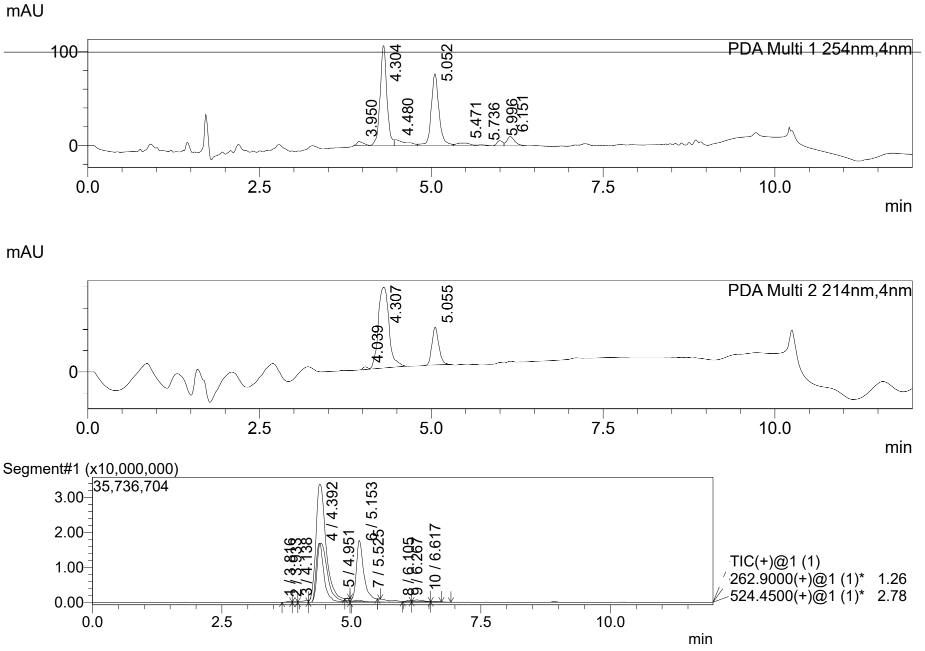


MS Spectrum

Peak#:1 R.Time:3.816(Scan#:353)
MassPeaks:11
Spectrum Mode:Averaged 3.803-3.824(352-354)
BG Mode:Calc Segment 1 - Event 1

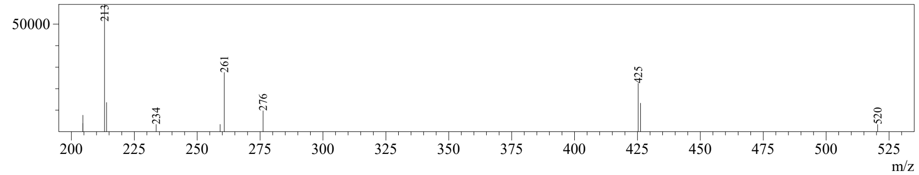

Figure S3 continued

**2700-015**



**<Sample Information>**

| | | | |
|---|---|---|---|
| Sample Name | : 2700 | | |
| Sample ID | : 2700-A15 peak 1 | | |
| Data Filename | : 2700-A15 peak 1 080521.lcd | | |
| Method Filename | : 6min 5-95, 1min clean(254)_(214) Col_1_LCMS3.lcm | | |
| Batch Filename | : 080521.lcb | | |
| Vial # | : 1-5 | Sample Type | : Unknown |
| Injection Volume | : 5 uL | | |
| Date Acquired | : 8/5/2021 4:06:42 PM | Acquired by | : System Administrator |
| Date Processed | : 9/20/2021 9:34:19 PM | Processed by | : System Administrator |

**<Chromatogram>**

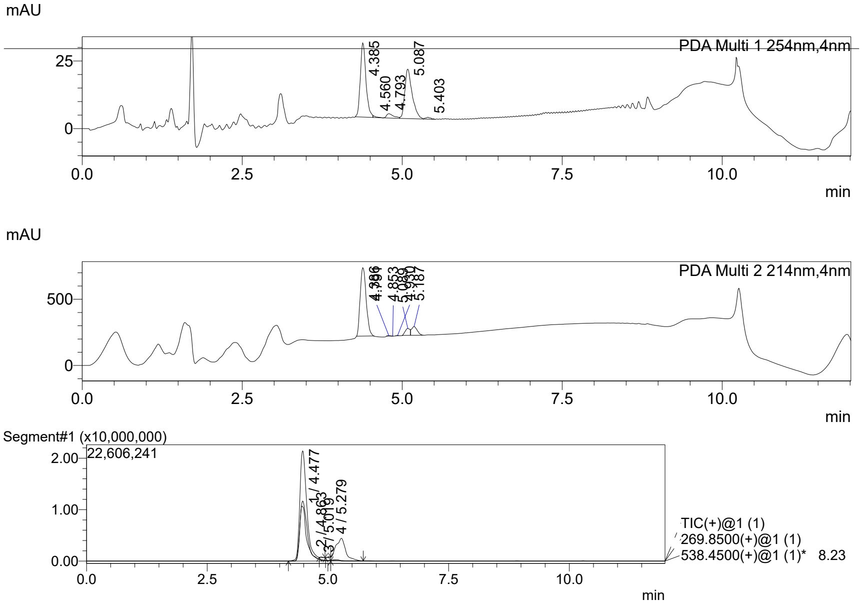


MS Spectrum

Peak#:1 R.Time:4.477(Scan#:414)
MassPeaks:64
Spectrum Mode:Averaged 4.463-4.485(413-415)
BG Mode:Calc Segment 1 - Event 1

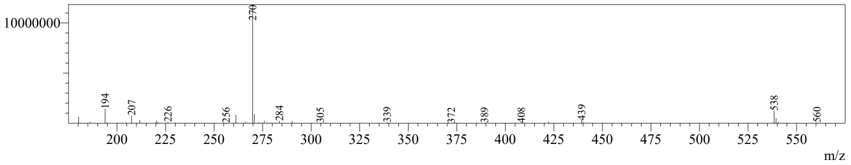




Figure S3. *Liquid chromatography-mass spectrometry (LCMS) analysis.* The figure shows UV traces at 254 nm, % purity integration data, Total ion current (TIC) data, mass spectrum (MS), and MS peak table.

Figure S4

*Checkerboard assays*

### 2700.001

| | Inhibition (%) Medians: Amk (µg/mL) | | | | | | Antimicrobial compound activity removed: Amk (µg/mL) | | | | | | AMK IC50/80: Effective Concentration | | AMK Fold Potentiation | |
|---|---|---|---|---|---|---|---|---|---|---|---|---|---|---|---|---|
| (µM) | 0 | 4 | 8 | 16 | 32 | 64 | 0 | 4 | 8 | 16 | 32 | 64 | 50% | 80% | 50% | 80% |
| 24.0 | 6.5% | 84.6% | 87.0% | 90.1% | 76.6% | | 0.0% | 83.6% | 86.1% | 89.4% | 74.9% | | 2.4 | 3.8 | 11.0 | 8.4 |
| 16.0 | 13.1% | 64.3% | 85.2% | 86.9% | 84.2% | | 0.0% | 58.9% | 83.0% | 84.9% | 81.8% | | 3.4 | 7.5 | 7.8 | 4.3 |
| 8.0 | 13.8% | 31.7% | 44.7% | 80.0% | 80.9% | | 0.0% | 20.8% | 35.9% | 76.7% | 77.8% | | 10.8 | > 32 | 2.5 | 1.0 |
| 4.0 | 14.2% | 20.8% | 29.8% | 47.1% | 75.9% | | 0.0% | 7.6% | 18.2% | 38.3% | 71.9% | | 21.6 | > 32 | 1.2 | 1.0 |
| 2.0 | 8.8% | 14.4% | 18.7% | 31.5% | 83.3% | | 0.0% | 6.2% | 10.9% | 24.9% | 81.7% | | 23.1 | 31.5 | 1.1 | 1.0 |
| 0.0 | 0.0% | 2.5% | 5.5% | 10.9% | 71.1% | | 0.0% | 2.5% | 5.5% | 10.9% | 71.1% | | 26.4 | > 32 | 1.0 | 1.0 |

### 2700.003

| | Inhibition (%) Medians: Amk (µg/mL) | | | | | | Antimicrobial compound activity removed: Amk (µg/mL) | | | | | | AMK IC50/80: Effective Concentration | | AMK Fold Potentiation | |
|---|---|---|---|---|---|---|---|---|---|---|---|---|---|---|---|---|
| (µM) | 0 | 4 | 8 | 16 | 32 | 64 | 0 | 4 | 8 | 16 | 32 | 64 | 50% | 80% | 50% | 80% |
| 24.0 | 1.4% | 89.2% | 90.1% | 84.9% | 88.0% | | 0.0% | 89.0% | 90.0% | 84.7% | 87.9% | | 2.2 | 3.6 | 10.9 | 8.9 |
| 16.0 | 11.1% | 73.7% | 85.9% | 89.5% | 90.0% | | 0.0% | 70.4% | 84.1% | 88.2% | 88.7% | | 2.8 | 6.8 | 8.6 | 4.7 |
| 8.0 | 11.0% | 33.4% | 43.3% | 78.0% | 83.0% | | 0.0% | 25.2% | 36.3% | 75.3% | 80.9% | | 10.8 | 29.3 | 2.3 | 1.1 |
| 4.0 | 11.9% | 21.7% | 27.1% | 41.3% | 54.8% | | 0.0% | 11.2% | 17.3% | 33.4% | 48.7% | | > 32 | > 32 | 1.0 | 1.0 |
| 2.0 | 8.6% | 14.8% | 17.4% | 32.2% | 15.8% | | 0.0% | 6.8% | 9.6% | 25.8% | 7.9% | | > 32 | > 32 | 1.0 | 1.0 |
| 0.0 | 0.0% | 2.5% | 9.8% | 22.6% | 25.2% | | 0.0% | 2.5% | 9.8% | 22.6% | 25.2% | | > 32 | > 32 | 1.0 | 1.0 |

### 2700.004

| | Inhibition (%) Medians: Amk (µg/mL) | | | | | | Antimicrobial compound activity removed: Amk (µg/mL) | | | | | | AMK IC50/80: Effective Concentration | | AMK Fold Potentiation | |
|---|---|---|---|---|---|---|---|---|---|---|---|---|---|---|---|---|
| (µM) | 0 | 4 | 8 | 16 | 32 | 64 | 0 | 4 | 8 | 16 | 32 | 64 | 50% | 80% | 50% | 80% |
| 24.0 | 10.6% | 21.0% | 26.5% | 86.6% | 84.6% | | 0.0% | 11.6% | 17.8% | 85.0% | 82.7% | | 11.8 | 15.4 | 2.7 | 2.1 |
| 16.0 | 9.7% | 15.8% | 18.5% | 39.3% | 88.2% | | 0.0% | 6.8% | 9.7% | 32.7% | 86.9% | | 21.1 | 30.0 | 1.5 | 1.1 |
| 8.0 | 11.1% | 14.7% | 16.4% | 26.2% | 78.0% | | 0.0% | 4.1% | 6.0% | 17.1% | 75.2% | | 25.1 | > 32 | 1.3 | 1.0 |
| 4.0 | 9.4% | 12.0% | 12.8% | 23.1% | 69.6% | | 0.0% | 2.8% | 3.7% | 15.0% | 66.4% | | 26.9 | > 32 | 1.2 | 1.0 |
| 2.0 | 0.0% | 1.1% | 4.0% | 26.4% | 87.6% | | 0.0% | 1.1% | 4.0% | 26.4% | 87.6% | | 22.2 | 30.0 | 1.4 | 1.1 |
| 0.0 | 0.0% | 4.0% | 8.0% | 16.0% | 32.0% | | 0.0% | 4.0% | 8.0% | 16.0% | 32.0% | | > 32 | > 32 | 1.0 | 1.0 |

### 2700.005

| | Inhibition (%) Medians: Amk (µg/mL) | | | | | | Antimicrobial compound activity removed: Amk (µg/mL) | | | | | | AMK IC50/80: Effective Concentration | | AMK Fold Potentiation | |
|---|---|---|---|---|---|---|---|---|---|---|---|---|---|---|---|---|
| (µM) | 0 | 4 | 8 | 16 | 32 | 64 | 0 | 4 | 8 | 16 | 32 | 64 | 50% | 80% | 50% | 80% |
| 24.0 | 3.4% | 13.8% | 84.0% | 86.5% | 86.5% | | 0.0% | 10.8% | 83.4% | 86.0% | 86.1% | | 6.2 | 7.8 | 4.0 | 4.1 |
| 16.0 | 6.5% | 12.8% | 22.2% | 89.7% | 88.6% | | 0.0% | 6.7% | 16.9% | 88.9% | 87.8% | | 11.7 | 15.0 | 2.1 | 2.1 |
| 8.0 | 7.7% | 11.8% | 15.0% | 63.1% | 85.1% | | 0.0% | 4.4% | 7.9% | 60.0% | 83.9% | | 14.5 | 29.4 | 1.7 | 1.1 |
| 4.0 | 8.1% | 11.9% | 14.2% | 28.9% | 82.8% | | 0.0% | 4.2% | 6.6% | 22.7% | 81.3% | | 23.5 | 31.7 | 1.0 | 1.0 |
| 2.0 | 5.8% | 9.6% | 10.5% | 24.2% | 75.9% | | 0.0% | 4.1% | 5.0% | 19.6% | 74.5% | | 24.9 | > 32 | 1.0 | 1.0 |
| 0.0 | 0.0% | 3.2% | 6.0% | 17.6% | 79.1% | | 0.0% | 3.2% | 6.0% | 17.6% | 79.1% | | 24.4 | > 32 | 1.0 | 1.0 |

### 2700.007

| | Inhibition (%) Medians: Amk (µg/mL) | | | | | | Antimicrobial compound activity removed: Amk (µg/mL) | | | | | | AMK IC50/80: Effective Concentration | | AMK Fold Potentiation | |
|---|---|---|---|---|---|---|---|---|---|---|---|---|---|---|---|---|
| (µM) | 0 | 4 | 8 | 16 | 32 | 64 | 0 | 4 | 8 | 16 | 32 | 64 | 50% | 80% | 50% | 80% |
| 24.0 | 8.5% | 15.8% | 23.2% | 38.8% | 87.8% | | 0.0% | 8.0% | 16.0% | 33.1% | 86.7% | | 21.0 | 30.0 | 1.2 | 1.1 |
| 16.0 | 11.3% | 16.4% | 20.3% | 29.5% | 84.9% | | 0.0% | 5.8% | 10.2% | 20.5% | 82.9% | | 23.6 | 31.2 | 1.0 | 1.0 |
| 8.0 | 8.5% | 14.4% | 16.3% | 26.0% | 84.1% | | 0.0% | 6.5% | 8.5% | 19.1% | 82.7% | | 23.8 | 31.3 | 1.0 | 1.0 |
| 4.0 | 7.2% | 13.0% | 15.6% | 29.4% | 88.5% | | 0.0% | 6.2% | 9.0% | 23.9% | 87.6% | | 22.6 | 30.1 | 1.1 | 1.1 |
| 2.0 | 4.6% | 11.4% | 15.3% | 29.0% | 89.8% | | 0.0% | 7.1% | 11.2% | 25.6% | 89.3% | | 22.1 | 29.7 | 1.1 | 1.1 |
| 0.0 | 0.0% | 5.6% | 11.4% | 40.2% | 87.8% | | 0.0% | 5.6% | 11.4% | 40.2% | 87.8% | | 19.3 | 29.4 | 1.3 | 1.1 |

### 2700.010

| | Inhibition (%) Medians: Amk (µg/mL) | | | | | | Antimicrobial compound activity removed: Amk (µg/mL) | | | | | | AMK IC50/80: Effective Concentration | | AMK Fold Potentiation | |
|---|---|---|---|---|---|---|---|---|---|---|---|---|---|---|---|---|
| (µM) | 0 | 4 | 8 | 16 | 32 | 64 | 0 | 4 | 8 | 16 | 32 | 64 | 50% | 80% | 50% | 80% |
| 24.0 | 8.2% | 23.4% | 85.6% | 90.2% | 90.3% | | 0.0% | 16.5% | 84.4% | 89.4% | 89.4% | | 6.0 | 7.7 | 4.1 | 4.1 |
| 16.0 | 21.0% | 22.1% | 25.2% | 88.5% | 89.8% | | 0.0% | 1.4% | 5.3% | 85.4% | 87.1% | | 12.5 | 15.5 | 2.0 | 2.1 |
| 8.0 | 18.4% | 19.5% | 21.4% | 25.2% | 48.2% | | 0.0% | 1.4% | 3.7% | 8.4% | 36.6% | | > 32 | > 32 | 1.0 | 1.0 |
| 4.0 | 17.3% | 21.2% | 20.1% | 25.0% | 40.3% | | 0.0% | 4.7% | 3.3% | 9.3% | 27.8% | | > 32 | > 32 | 1.0 | 1.0 |
| 2.0 | 14.0% | 17.8% | 18.8% | 23.1% | 37.9% | | 0.0% | 4.5% | 5.6% | 10.6% | 27.8% | | > 32 | > 32 | 1.0 | 1.0 |
| 0.0 | 0.0% | 4.9% | 9.3% | 22.9% | 81.9% | | 0.0% | 4.9% | 9.3% | 22.9% | 81.9% | | 23.4 | 31.5 | 1.0 | 1.0 |

### 2700.013

| | Inhibition (%) Medians: Amk (µg/mL) | | | | | | Antimicrobial compound activity removed: Amk (µg/mL) | | | | | | AMK IC50/80: Effective Concentration | | AMK Fold Potentiation | |
|---|---|---|---|---|---|---|---|---|---|---|---|---|---|---|---|---|
| (µM) | 0 | 4 | 8 | 16 | 32 | 64 | 0 | 4 | 8 | 16 | 32 | 64 | 50% | 80% | 50% | 80% |
| 24.0 | -5.6% | 12.0% | 22.9% | 52.3% | 80.6% | | 0.0% | 16.6% | 27.0% | 54.8% | 81.7% | | 14.6 | 31.0 | 1.7 | 1.0 |
| 16.0 | 9.1% | 17.3% | 22.6% | 27.6% | 32.1% | | 0.0% | 9.1% | 14.9% | 20.4% | 25.4% | | > 32 | > 32 | 1.0 | 1.0 |
| 8.0 | 16.4% | 20.4% | 21.5% | 23.3% | 29.5% | | 0.0% | 4.8% | 6.1% | 8.3% | 15.7% | | > 32 | > 32 | 1.0 | 1.0 |
| 4.0 | 15.4% | 17.8% | 20.2% | 23.8% | 25.4% | | 0.0% | 2.9% | 5.7% | 10.0% | 11.9% | | > 32 | > 32 | 1.0 | 1.0 |
| 2.0 | 12.0% | 14.5% | 18.1% | 21.0% | 24.5% | | 0.0% | 2.9% | 7.0% | 10.2% | 14.2% | | > 32 | > 32 | 1.0 | 1.0 |
| 0.0 | 0.0% | 5.0% | 4.9% | 20.9% | 22.9% | | 0.0% | 5.0% | 4.9% | 20.9% | 22.9% | | > 32 | > 32 | 1.0 | 1.0 |

#### EC 2/3-Fold IC50/80

| Compound ID | 2-Fold 50% | 2-Fold 80% | 3-Fold 50% | 3-Fold 80% |
|---|---|---|---|---|
| 2700.001 | 6.5 | 10.4 | 8.8 | 12.9 |
| 2700.003 | 7.2 | 10.0 | 8.9 | 12.2 |
| 2700.004 | 19.3 | 23.4 | >24 uM | >24 uM |
| 2700.005 | 14.2 | 15.0 | 19.9 | 19.5 |
| 2700.007 | >24 uM | >24 uM | >24 uM | >24 uM |
| 2700.010 | 16.2 | 15.5 | 19.9 | 19.6 |
| 2700.013 | >24 uM | >24 uM | >24 uM | >24 uM |

Figure S4. *Checkerboard assays*. The leftmost column shows the measured values (medians %, n ≥ 6). The adjacent column shows the adjusted values after removing any antimicrobial activity exerted by the testing compound alone. Adjustment was carried out as described in Materials and Methods. The third column shows the amikacin inhibitory concentration 50 and 80% and the fold potentiation at the corresponding concentrations of compound. The fold potentiation was calculated dividing the IC50 and IC80 values in the absence of compound by the values obtained in the presence of each compound concentration. In those cases where the IC50 or IC80 values were >32, the value used for the calculation was 32. The rightmost column indicates the compound concentration needed for 2- or 3-fold potentiation of both the 50% and 80% inhibitory dose points.

Figure S5

*Molecular docking*

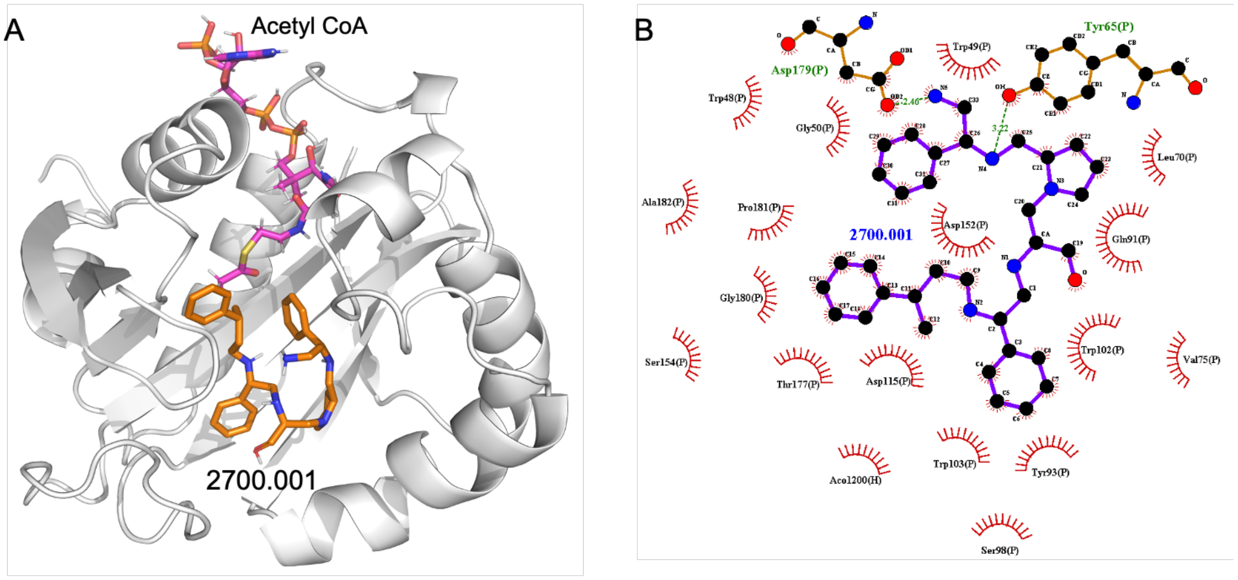


Figure S5a. *Complex between compound 2700.001 and AAC(6′)-Ib.* (A) The complex of 2700.001 and AAC(6′)-Ib obtained from molecular docking. The bound acetyl CoA is also shown. (B) Interaction map of the ligand in its binding site of the AAC(6′)-Ib receptor. The map shows that the phenol group of Tyr65 forms a hydrogen bond with 2700.001. The primary amine of 2700.001 forms a hydrogen bond with Asp179. The pictures show the top-ranked, lowest energy conformation.

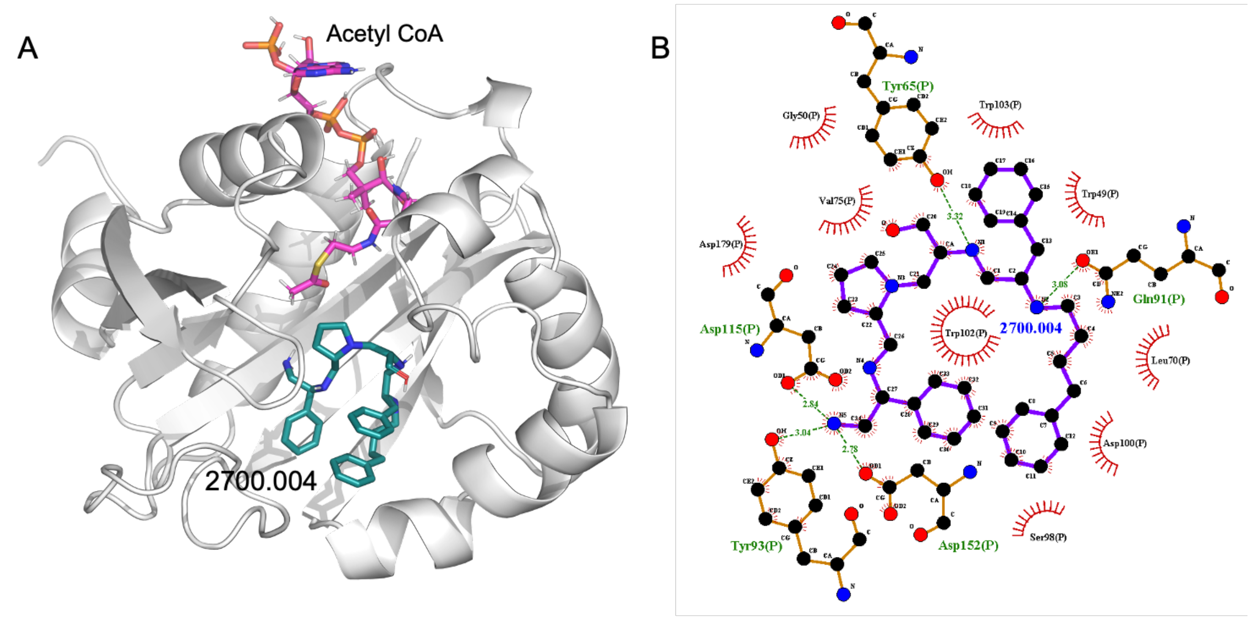


Figure S5b. *Complex between compound 2700.004 and AAC(6′)-Ib.* (A) The complex of 2700.004 and AAC(6′)-Ib obtained from molecular docking. The bound acetyl CoA is also shown. The figure shows a potential for intramolecular pi stacking between two phenol groups. (B) Interaction map of the ligand in its binding site of the AAC(6′)-Ib receptor. The map shows that 2700.004 forms several bonds with the receptor, particularly the primary amine, which forms hydrogen bonds with Asp115, Asp152, and the phenol group of Tyr93. Gln91 and the phenol group of Tyr65 also form hydrogen bonds with the other side of the molecule.

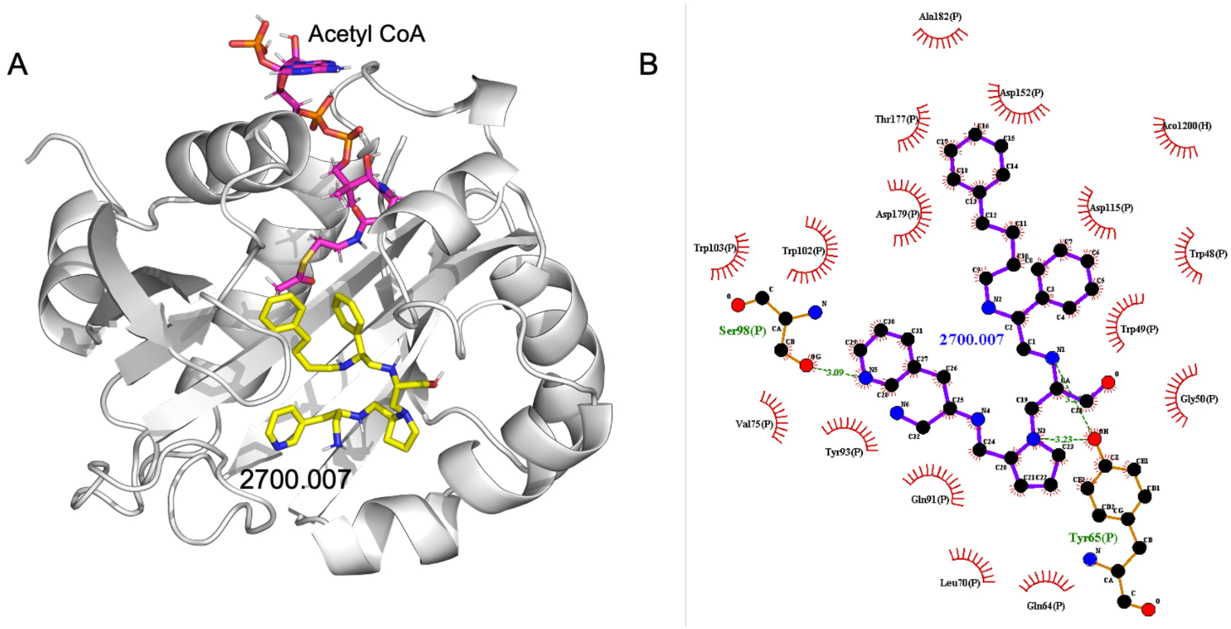


Figure S5c. *Complex between compound 2700.007 and AAC(6′)-Ib.* (A) The complex of 2700.007 and AAC(6′)-Ib obtained from molecular docking. The bound acetyl CoA is also shown. (B) Interaction map of the ligand in its binding site of the AAC(6′)-Ib receptor. The map shows that although there is conformational change, 2700.007 maintains a hydrogen bond with the phenol group of Tyr65 and hydrogen bonding with Ser98.

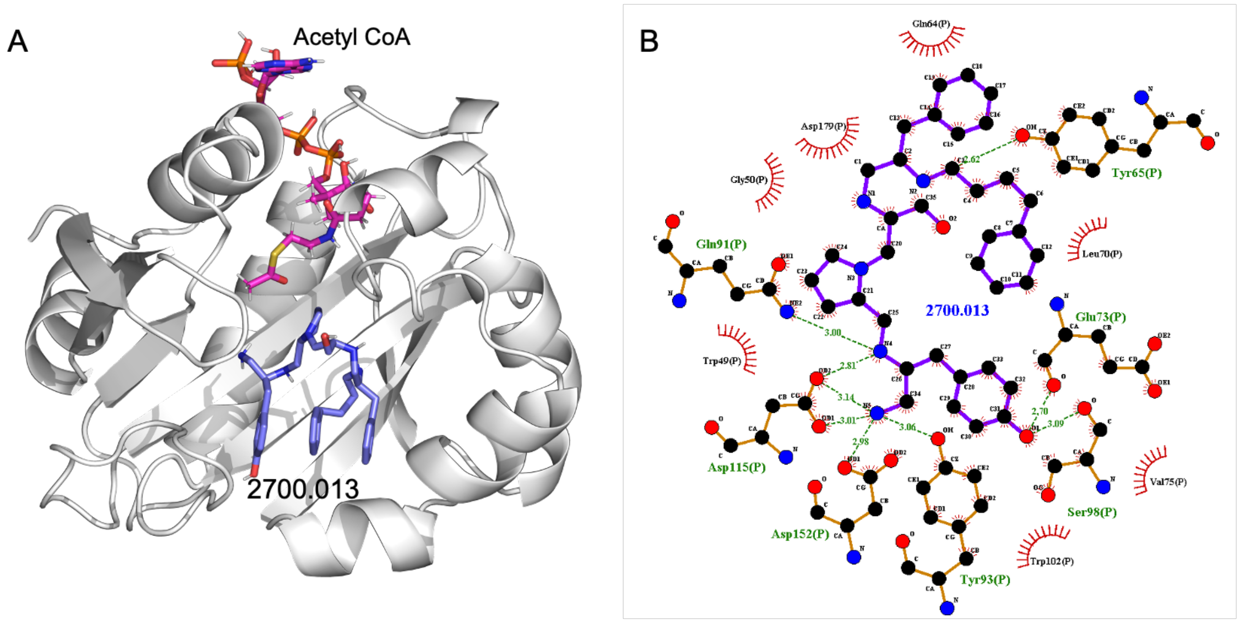


Figure S5d. *Complex between compound 2700.013 and AAC(6′)-Ib.* (A) The complex of 2700.013 and AAC(6′)-Ib obtained from molecular docking. The bound acetyl CoA is also shown. (B) Interaction map of the ligand in its binding site of the AAC(6′)-Ib receptor. The map shows 2700.013 forms hydrogen bonds through its primary amine with Tyr 93, Asp115, and Asp152. Sr98 and Glu73 shows hydroxyl group functionality while the Ser98 and the phenol group of Tyr65 show hydrogen bonding with 2700.013.
